# *Stenotrophomonas maltophilia* exhibits defensive multicellularity in response to a *Pseudomonas aeruginosa* quorum sensing molecule

**DOI:** 10.1101/2025.05.02.651457

**Authors:** Stefan Katharios-Lanwermeyer, Tiffany M. Zarrella, Marshall Godsil, Stefanie Severin, Alejandro E. Casiano, Chin-Hsien Tai, Anupama Khare

## Abstract

Microorganisms commonly exist in polymicrobial communities, where they can respond to interspecies secreted molecules by altering behaviors and physiology, however, the underlying mechanisms remain underexplored. Here we investigated interactions between *Stenotrophomonas maltophilia* and *Pseudomonas aeruginosa*, co-infecting opportunistic pathogens found in pneumonia and chronic lung infections, including in cystic fibrosis. We found that *S. maltophilia* forms robust protective multicellular aggregates upon exposure to *P. aeruginosa* secreted factors. Experimental evolution for lack of aggregation selected for fimbrial mutations and we found that fimbriae are required on both interacting *S. maltophilia* cells for aggregation. Untargeted metabolomics and targeted validations revealed that the quorum sensing molecule *Pseudomonas* quinolone signal (PQS) directly induced *S. maltophilia* aggregation, and co-localized with the aggregates. Further, in co-culture with *P. aeruginosa*, wild-type *S. maltophilia* formed aggregates, resulting in up to 75-fold increased survival from *P. aeruginosa* competition compared to fimbrial mutants. Finally, multiple other bacterial species similarly aggregated upon exposure to *P. aeruginosa* exoproducts, indicating a more general response. Collectively, our work identifies a novel multispecies interaction where a quorum sensing molecule from a co-infecting pathogen is sensed as a ‘danger’ signal, thereby inducing a protective multicellular response.

## INTRODUCTION

Polymicrobial communities, where two or more microbial species share an environmental niche, predominate in diverse settings including chronic airway infections, wounds, indwelling catheters, and natural soil and aquatic environments (*1–6*). Within mixed-species settings, microbes participate in cooperative and competitive interactions via metabolic crosstalk, nutrient sequestration, contact-dependent inhibition and production of antimicrobials (*5*, *7–11*). In addition to directly altering microbial growth and survival, interspecies interactions can also lead to behavioral changes, such as emergence of motility or biofilm formation (*12–14*). These behaviors include responses to danger sensing, where bacteria sense cues, often resulting from nearby cellular damage, as imminent threats and initiate defensive or antagonistic responses (*15–18*). Bacteria may also engage in competition sensing, where nutrient depletion and other stresses can result in competitive responses (*19*).

*Pseudomonas aeruginosa* is an opportunistic pathogen, that is prevalent in diverse polymicrobial clinical and environmental niches where it interacts with a variety of cohabiting species (*20–22*). One such organism is *Stenotrophomonas maltophilia*, a gram-negative emerging opportunistic pathogen intrinsically resistant to multiple antibiotics (*23*). *P. aeruginosa* and *S. maltophilia* are found together in clinical multispecies environments like chronic airway infections in people with cystic fibrosis (*24*, *25*) where they are associated with accelerated lung decline (*26*, *27*) and in non-CF pneumonias where co-isolation with *P*. *aeruginosa* is associated with elevated mortality (*28*). They may also co-occur in polymicrobial chronic wounds and burn-associated infections (*29–31*).

Despite the ubiquity and clinical importance of *P. aeruginosa* in the CF airway, its interactions with emergent members of the CF polymicrobial community, such as *S. maltophilia*, are only beginning to be understood. *P. aeruginosa* increases persistence of *S. maltophilia* in murine lung infections (*32*), and enhances *S. maltophilia* adherence to respiratory epithelial cells in culture, putatively mediated by the type IV pilus (*33*). Antagonistic interactions have also been observed, where *S. maltophilia* can inhibit *P. aeruginosa* fitness in a contact-dependent manner via both the type VI secretion system (T6SS) (*34*) and type IVa secretion system (T4SS) (*35*). Conversely, *P. aeruginosa* also exhibits contact-dependent growth inhibition of *S. maltophilia*, while *S. maltophilia* can alter *P. aeruginosa* biofilm architecture, and both species may alter motility in the other (*36–38*).

The *P. aeruginosa* secretome is complex and has been extensively investigated (*39–41*), revealing evidence of contact-independent interspecies antagonism against competing organisms. For example, the *<u>P</u>seudomonas* <u>q</u>uinolone <u>s</u>ignal (PQS) can inhibit *Aspergillus fumigatus* growth and biofilm formation through iron sequestration (*42*). Additionally, *P. aeruginosa* can repress *Staphylococcus aureus* cellular respiration through the production of the antimicrobial pigment pyocyanin and the alkyl quinolone HQNO (*43*, *44*), molecules that also reduce growth and viability, similar to other secreted products like proteases, and iron sequestering siderophore molecules (*44–46*). *P. aeruginosa* can induce changes in behavior with the secreted surfactant rhamnolipids inducing dispersion of *Desulfovibrio vulgaris* and *Escherichia coli* biofilms (*22*). However, the role of *P. aeruginosa* secreted factors in interactions with *S. maltophilia* remains poorly understood.

In this study we investigate whether *S. maltophilia* responds to *P. aeruginosa*-produced secreted factors, examining the nature of this response, and the underlying molecules and mechanisms. Using genetics, biochemistry, and microscopy, we identify an emergent multicellular defensive behavior of *S. maltophilia* in response to *P. aeruginosa* secreted factors. We find that PQS is necessary and sufficient to induce *S. maltophilia* aggregation. Further, we demonstrate that *S. maltophilia* aggregation is mediated by Smf-1 fimbriae and provides protection from *P. aeruginosa* in co-culture. Our data thus reveal that *S. maltophilia* senses *P. aeruginosa* PQS as a danger signal and mounts a protective behavioral response. Finally, we provide evidence that transition to multicellularity triggered by *P. aeruginosa* is a more general response also observed in additional bacterial species.

## RESULTS

### *S. maltophilia* aggregates in response to interspecies metabolites produced by *P. aeruginosa*

Given the existence of *S. maltophilia* (*Sm*) and *P. aeruginosa* (*Pa*) in similar environmental niches and in clinical co-infections, we hypothesized that exposure to *Pa* secreted metabolites could directly antagonize or alter *Sm* behavior and physiology. To test this, we assessed the impact of 50% (v/v) *Pa* cell-free supernatant on *Sm* culture growth over time, compared to supernatant derived from *S. aureus* and *Sm* itself, as well as an M63 medium salts control. Only the addition of *Pa* supernatant to shaken *Sm* cultures resulted in an initial 3-log decrease in *Sm* colony-forming units (CFUs) per mL (**Figure 1A**). However, between 4 and 8 h, the population started to recover, and eventually reached levels seen in the other conditions by the terminal 24-hour time point.

**Figure 1.**
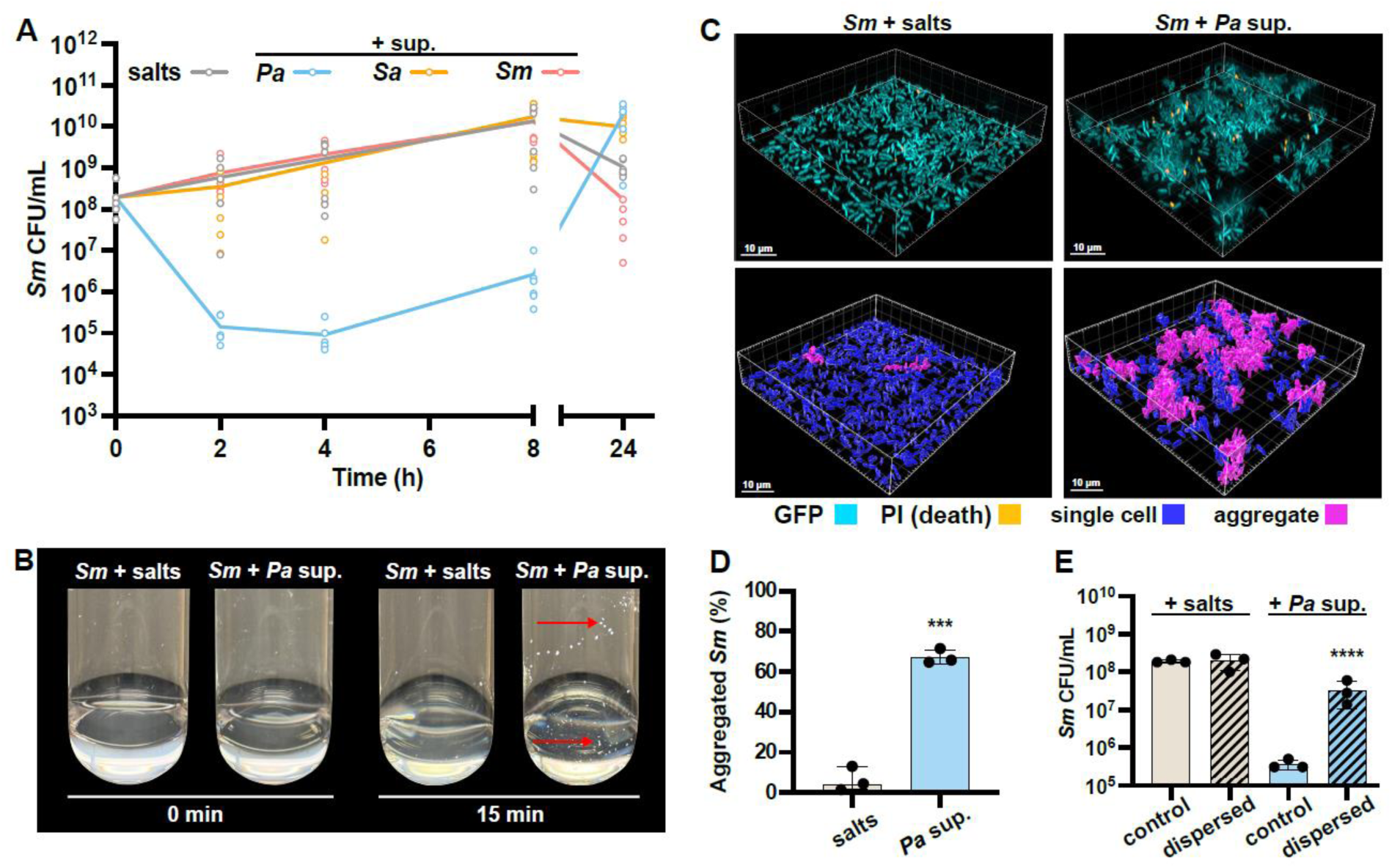
*Sm* aggregates in response to *Pa* supernatant. **(A)** *S. maltophilia* (*Sm*) was exposed to cell-free supernatant derived from *P. aeruginosa* (*Pa*), *Staphylococcus aureus* (*Sa*), or *Sm* and colony forming units (CFUs) assessed from 2-24 h. **(B)** Tubes of *Sm* cells exposed to salts and *Pa* supernatant, and imaged before and after 15 min of shaking incubation. Red arrows indicate some of the visible macroscopic aggregates. **(C)** Fluorescence microscopy was employed to image GFP-tagged *Sm* that were stained with propidium iodide (PI) in a static 96-well plate and imaged after 2 h exposure to salts or *Pa* supernatant. (Top) GFP and PI fluorescence is shown. (Bottom) Aggregates, defined as any object with a biovolume exceeding 20 μm^3^, are pseudo-colored in pink while non-aggregates are in blue. **(D)** The percent biovolume of *S. maltophilia* cells present within aggregates upon exposure to salts or *Pa* supernatant. **(E)** CFU enumeration of *Sm* after 2 h exposure to salts or *Pa* supernatant followed by mechanical dispersion. **(B, C)** Representative images of three biological replicates are shown. **(A, D, E)** Data shown are **(A)** mean and all individual replicates or **(D, E)** the mean ± SD for three biological replicates. Significance is shown for comparison to the salts control, as tested by **(A)** a two-way ANOVA with the Dunnett’s test for multiple comparisons **(D)** an unpaired *t*-test, and **(E)** a one-way ANOVA on log-transformed values followed by the Tukey’s test for multiple comparisons (***, *p* < 0.001; ****, *p* < 0.0001).

We observed that *Sm* formed macroscopic clumps after as little as 15 minutes exposure to *Pa* supernatant (**Figure 1B**). We reasoned that the decreased CFUs we observed may not reflect cell death but instead the exclusion of bacterial clumps from sampling, or sampled multicellular aggregates forming single colonies, thus reducing the apparent CFU/mL. Subsequent increase in cell density may result from growth of single, non-aggregated cells.

To directly test for the formation of bacterial aggregates as well as cell death, we performed fluorescent microscopy and live/dead imaging on static *Sm* cultures exposed to either the salts control or *Pa* supernatant for 2 h in glass-bottom 96-well dishes. Staining with propidium iodide (PI), a membrane-impermeant dye that only stains membrane-compromised cells (*47*) validated that there was no significant increase in *Sm* cell death upon exposure to *Pa* supernatant compared to the control (**Figure 1C** and **Supplemental Figure 1**). Additionally, *Sm* formed large, three-dimensional aggregates in the presence of *Pa* secreted products (**Figure 1C**). We quantified aggregation using single-cell segmentation and thresholding, defining any bacterial object exceeding 20 μm³ as an aggregate. Quantification of the biovolume of single cells versus multicellular aggregates confirmed that single cells predominated in the salts control, whereas aggregates accounted for the majority (∼66%) of *Sm* biovolume when exposed to *Pa* exoproducts (**Figure 1D**). To test our hypothesis that the decrease in *Sm* CFUs in response to *Pa* secreted factors (**Figure 1A**) was caused by aggregation, samples were detached via vortexing, and mechanically dispersed prior to CFU enumeration. We reasoned that dispersion of bacterial aggregates would liberate single cells which would form individual colonies, thereby providing a more accurate readout of live cell numbers. Consistent with this, dispersion rescued CFUs of *Sm* aggregates induced by *Pa* supernatant but had no impact on non-aggregated *Sm* treated with the salts control (**Figure 1E**). Finally, we found that *Pa* supernatant-induced *Sm* aggregation was a dose-dependent response. *Sm* aggregation was absent with 5% *Pa* supernatant, but readily occurred with 12% supernatant, and *Sm* aggregate biovolume subsequently increased up to 50% (v/v) *Pa* supernatant (**Supplemental Figure 2**). Collectively, these findings provide evidence that *Sm* engages in rapid and robust bacterial aggregation in response to *Pa* secreted factors, a phenotype that can be accurately assayed through cellular enumeration where aggregation results in a decrease in *Sm* CFUs, and microscopically via biovolumetric analysis.

### *S. maltophilia* aggregation in response to *P. aeruginosa* metabolites is an active behavior and is observed in clinical isolates

Bacterial aggregation is likely a ubiquitous mode of multicellular organization, observed among species as varied as *E. coli* and *S. aureus* (*48*, *49*), and in niches as diverse as aquatic environments and the gastrointestinal tract (*50–52*). *Pa* has been shown to aggregate in response to environmental polymers, such as extracellular DNA, mucin, and polyethylene, a behavior that was maintained even in UV-killed bacteria (*53*). We therefore tested whether *Sm* aggregation in response to *Pa* supernatant similarly occurs among inert and dead cells, or is an active process requiring respiration, ATP generation, or protein synthesis. We found that upon *Sm* incubation under hypoxic conditions, subsequent exposure to *Pa* supernatant failed to induce aggregation and instead resulted in elevated cell death as measured by PI staining (**Supplemental Figure 3A, C, D**). Similarly, *Sm* pre-treatment with carbonyl cyanide-m-chlorophenylhydrazone (CCCP), which disrupts the proton motive force (*54*), also led to cell death and a lack of *Sm* aggregation in response to *Pa* supernatant (**Supplemental Figure 3B, E, F**). Thus, while *Pa* supernatant is not significantly toxic to *Sm* by itself, the combinatorial impact of *Pa* metabolites and hypoxia or ATP depletion can result in reduced *Sm* cell viability.

In contrast, pre-treatment with the antibiotic chloramphenicol, which inhibits protein synthesis, prior to exposure to *Pa* supernatant, resulted in a significant reduction of *Sm* aggregation without an increase in cell death (**Supplemental Figure 3B, E, F**). These findings provide evidence that robust *Sm* aggregation in response to *Pa* secreted factors is an active process that requires protein synthesis. Further, hypoxia, ATP depletion, or inhibition of protein synthesis did not lead to *Sm* aggregation in the absence of *Pa* supernatant indicating that aggregation is not an *Sm* response to these stresses.

We next sought to understand whether this interaction is more broadly observed in clinical CF isolates of *Sm* and *Pa*. Upon exposure to supernatant from our reference *Pa* strain PA14, two of the four *Sm* isolates we tested (CF077 and CF082) aggregated, while the other two (CF087 and CF091) did not (**Figure 2A, 2B**). Similarly, two out of four *Pa* clinical isolates (CF033 and *Pa* CF081) induced aggregation in our reference *Sm* strain K279a similar to the *Pa* PA14 levels, a third one (CF047) induced a lower level of aggregation, while the fourth one (CF002) did not induce *Sm* aggregation (**Figure 2C, 2D**). Cumulatively, these results suggest that interspecies aggregation of *Sm* induced by *Pa* secreted factors occurs among select clinical isolates of both species.

**Figure 2.**
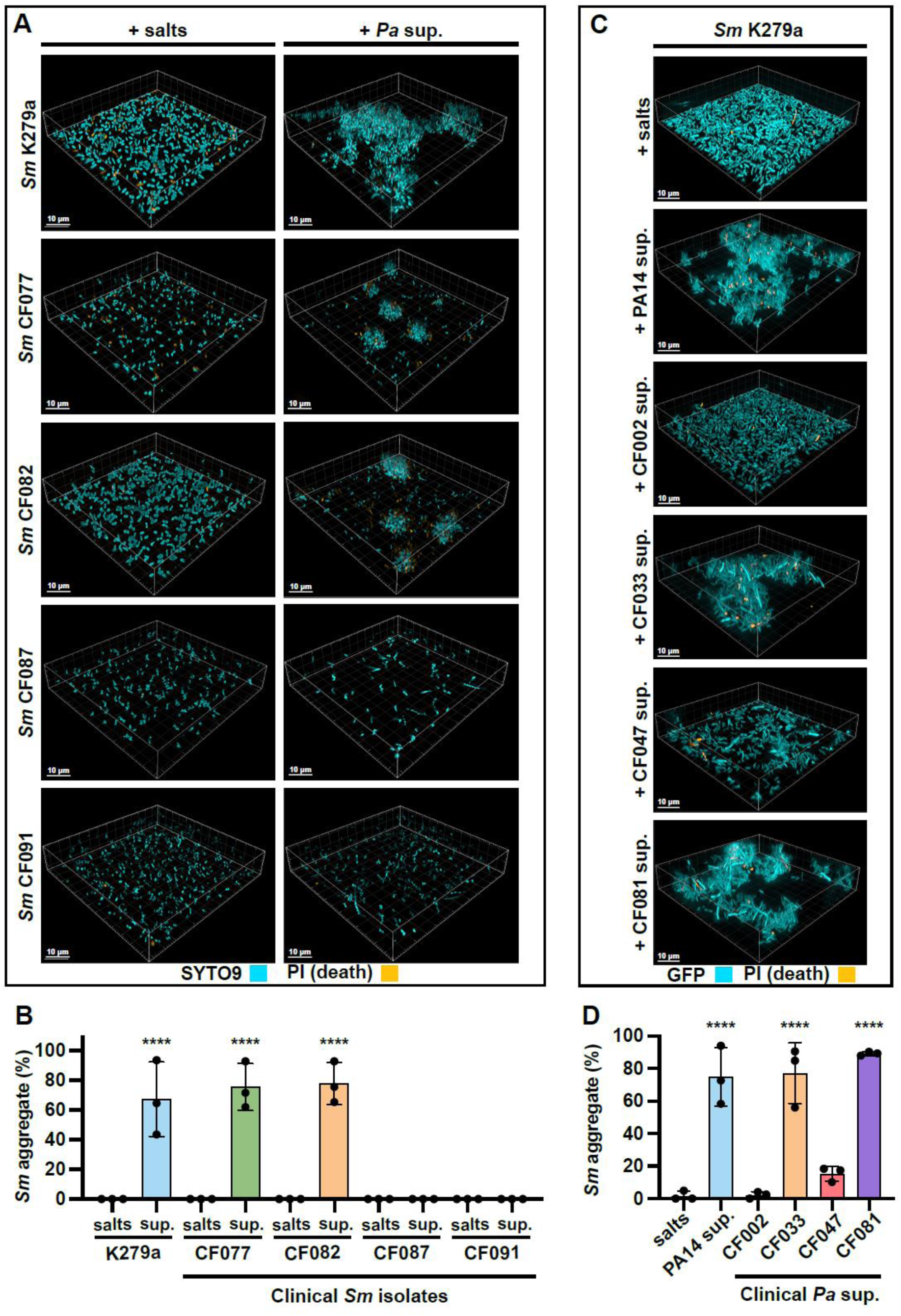
Select CF isolates from both species engage in interspecies aggregation. **(A, B)** *Sm* K279a reference strain and CF clinical isolates were stained with SYTO9 and propidium iodide and aggregation in response to PA14 cell-free supernatant analyzed via **(A)** microscopy and **(B)** biovolume quantification. **(C, D)** *Pa* supernatant was derived from *Pa* reference strain PA14 and CF clinical isolates and aggregation induction of *Sm* K279a assessed via **(C)** microscopy and **(D)** biovolume quantification. **(A, C)** Representative images of three biological replicates are shown. **(B, D)** Data shown are the mean ± SD for three biological replicates. Significance is shown for comparison to the respective salts control, as tested by a one-way ANOVA followed by the Tukey’s test for multiple comparisons (****, *p* < 0.0001).

### *Sm* Smf-1 fimbriae are required for aggregation in response to *Pa* secreted factors and environmental mucins

To identify the genetic requirements for *Sm* aggregation in response to *Pa* secreted factors, we used experimental evolution, reasoning that after exposure to *Pa* supernatant, the planktonic fraction would be enriched for cells that did not aggregate. We thus exposed *Sm* to *Pa* supernatant for ∼20 hours and then used an aliquot of cells from the planktonic fraction as the inoculum for the next round of *Pa* supernatant exposure, thereby selecting for aggregation-deficient strains. We repeated such passaging 5 and 10 times for two independently evolved populations.

Whole-genome sequencing (WGS) of isolates from these populations revealed that each had mutations upstream of or in the coding sequence of *smf-1* (*RS03355*), a gene encoding the subunit of the Smf-1 fimbriae (**Figure 3A and Supp. Table 1**), the only gene that showed mutations in all sequenced isolates. The Smf-1 fimbriae have been previously implicated in biofilm formation and epithelial cell attachment (*55*, *56*), but a role in cell-cell aggregation has not been reported. Examination of the locus immediately downstream of *smf-1* identified a putative four-gene operon with three genes of unknown function—*RS03360*, *RS03365*, and *RS03370*—which are annotated as a putative chaperone, outer-membrane usher protein, and fimbrial protein, respectively. The Smf-1 fimbria is thus likely a chaperone-usher fimbria (*57*, *58*) encoded by this operon, and we named the three downstream genes *smfB*, *smfC*, and *smfD*, respectively. To better understand *SmfD* function, we identified homologous proteins. The top-scoring BLASTP match for SmfD was MrkD, an adhesin required for collagen attachment in *Klebsiella pneumoniae* (*59*, *60*), with 33% identity and 49% similarity and other known adhesins also showed homology, indicating that SmfD likely encodes an adhesin.

**Figure 3.**
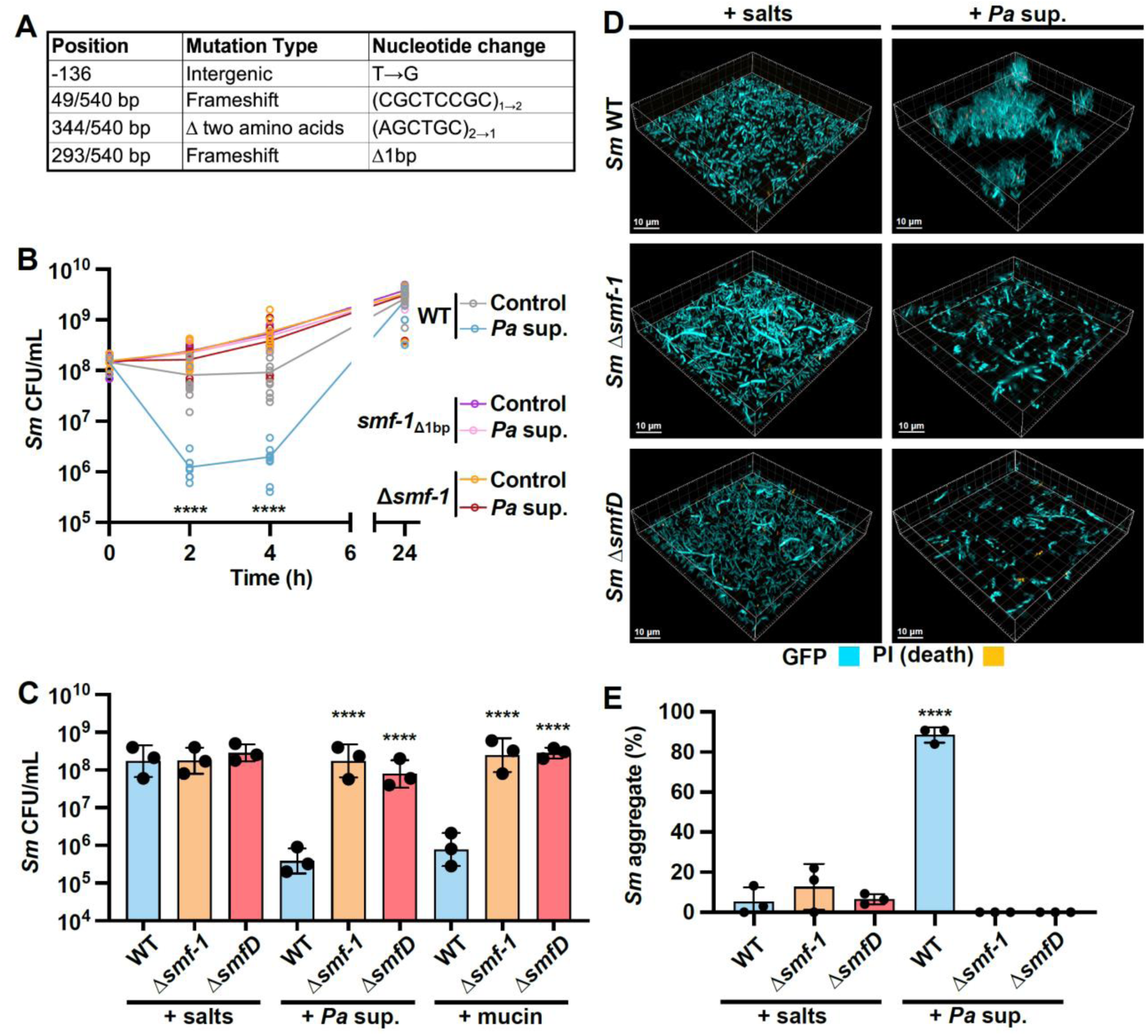
Smf-1 fimbriae are required for *Sm* aggregation in response to *Pa* CFS. **(A)** Position and sequence alteration of mutations in the 540 bp gene *smf-1* acquired after selection of aggregate-negative isolates during serial *Pa* cell-free supernatant exposure. **(B)** CFU enumeration of *Sm* WT, *smf-1* single base pair deletion (*smf-*1_Δ1bp_) and Δ*smf-1* clean deletion upon exposure to *Pa* supernatant. **(C-E)** WT, *Δsmf-1*, and Δ*smfD* exposed to either the salts control, *Pa* supernatant, or mucin, for 2 h, and aggregation analyzed via **(C)** CFU enumeration, **(D)** microscopy, or **(E)** the percentage of biovolume present in aggregates. **(B, C, E)** Data shown are for three biological replicates and represented as **(B)** the mean and all replicates or **(C, E)** mean ± SD. Significance is shown for comparison to **(B, E)** the respective salts control and **(C)** the respective WT control as tested by a **(B)** two-way ANOVA followed by the Tukey’s test for multiple comparisons, and **(C, E)** one-way ANOVA on **(C)** log-transformed or **(E)** measured values followed by the Tukey’s test for multiple comparisons (****, *p* < 0.0001). **(D)** Representative images of three biological replicates are shown.

We next determined additional responses of *Sm* to *Pa* secreted factors using RNA-seq to compare the transcriptional profile of WT *Sm* exposed to *Pa* secreted products versus the salts control. After 30 minutes of challenge with *Pa* supernatant, we observed upregulation of genes involved in iron homeostasis, transport, and aerobic respiration, and downregulation of flagellar- and Type IV pili-based motility, as well as anaerobic respiration (**Supplemental Table 2**), indicating that *Pa* supernatant likely leads to iron competition, and alters *Sm* metabolism and motility. The *smfB*, *smfC*, and *smfD* genes were also upregulated upon exposure to *Pa* supernatant, while *smf-1* trended towards a significant increase (**Supplemental Figure 4**), further implicating the Smf fimbriae in the *Sm* response.

To determine if Smf-1 fimbriae are required for *Sm* aggregation in response to *Pa* supernatant, we introduced the 1 bp deletion we identified in *smf-1* through experimental evolution, and clean deletions lacking the entire coding region of either *smf-1* or *smfD*, into the parental background. We first observed that the *smf-1*_Δ1bp_, and Δ*smf-1* strains did not show the decrease in CFU per mL in response to *Pa* supernatant that was seen for the wild-type (WT), indicating a deficiency in aggregation (**Figure 3B**). A previous report had shown that *Sm* K279a formed large, multicellular structures when grown in the artificial sputum medium SCFM2, likely due to the glycoprotein polymer mucin in the medium (*52*, *61*). We therefore also tested aggregation in the presence of mucin and found that unlike the WT, the Δ*smf-1* and Δ*smfD* mutants were aggregation-negative when challenged with either *Pa* supernatant or mucin, as measured by CFU enumeration (**Figure 3C**). Lack of aggregation in these mutants upon exposure to *Pa* metabolites was also observed via microscopy (**Figure 3D, E**).

We also tested if Smf-1 is required for bridge interactions between adjacent dells. While a 1:1 mixture of WT cells with different fluorophores (GFP and mKO) formed mixed aggregates upon exposure to *Pa* supernatant, under the same conditions Δ*smf-1* mKO cells did not co-aggregate with WT GFP cells (**Supplemental Figure 5A**). These findings support the view that *Sm* aggregation in response to *Pa* secreted factors occurs through a Smf-1 fimbriae-mediated bridging interaction between different cells.

Through the course of imaging Δ*smf-1* and Δ*smfD*, we noticed that both mutants exhibited enhanced motility in addition to being aggregation-negative. This is evident in **Figure 3D**, where the apparent chains of cells seen for the Δ*smf-1* and Δ*smfD* mutants actually show individual cells rapidly moving in the culture and being imaged in multiple Z-stacks. To test if the inability to aggregate in response to *Pa* supernatant was simply a function of this enhanced motility, we introduced a Δ*fliI* mutation lacking a putative flagellum-specific ATPase required for *Sm* swimming motility (*62*) into both the WT and Δ*smf-1* backgrounds. Agar-based swim assays confirmed the enhanced motility of Δ*smf-1*, which was abolished in the Δ*smf-1* Δ*fliI* double mutant (**Supplemental Figure 5B**). Further, we observed no change in aggregation ability between Δ*smf-1* and Δ*smf-1* Δ*fliI* (**Supplemental Figure 5C**), which suggested that loss of aggregation by Δ*smf-1* was specifically due to the absence of the *smf-1* subunit and not due to changes in motility.

To further validate that the loss of *Sm* aggregation in response to *Pa* supernatant was due to the absence of the Smf-1 fimbriae, we complemented the Δ*smf-1* mutant with the pBBR1MCS plasmid (*63*) containing the entire *smf* operon (pSmf). Upon exposure to *Pa* secreted products we observed an almost complete recovery of *Pa* supernatant-induced aggregation in Δ*smf-1* complemented with pSmf, as measured by microscopy (**Supplemental Figure 5D, E**), and a partial rescue of *Sm* aggregation as measured by a ∼1-log decrease in CFU per mL (**Supplemental Figure 5F**). These results demonstrate that Smf-1 is required for *Sm* aggregation induced by *Pa* supernatant but indicates that the amount of Smf-1 expressed may alter aggregate integrity. The discrepancy we observed between microscopy versus CFU-based assays may indicate that pSmf aggregates are less robust to withstanding agitation and dilution employed during CFU enumeration and thus do not lead to a drastic reduction in CFU.

To better understand the interaction between Smf-1 fimbriae and the SmfD protein, we predicted the structures of the complex using AlphaFold (**Supplemental Figure 6A, B**). Similar to *E. coli* FimA fimbriae (*58*), *Sm* Smf-1 is predicted to interact with SmfD as well as Smf-1 subunits via donor strand complementation. In particular, Smf-1 residues 24-33 formed an extended β-strand that inserts between SmfD residues 206-216 and 330-339 to complete the β-barrel structure. Likewise, the β-strand of the second Smf-1 residue 24-33 (**Supplemental Figure 6B, C**) inserts between the two β-strands 45–53 and 166-175 of the first Smf-1 to complete the beta-barrel.

We also examined the domain structure of SmfD using UniProt (*64*), and identified a MrkD-like receptor-binding domain (RBD) in residues 41-187. Subsequent DELTA-BLAST analysis of this SmfD RBD revealed high homology with both MrkD and Abp1D, an adhesin in *Acinetobacter baumannii* involved in fibrinogen binding (*65*) (**Supplemental Figure 6D**), and conservation of *K. pneumoniae* MrkD residues required for collagen binding (*66*). Two such residues, V91 and R102, are part of a hydrophobic binding patch in MrkD (*66*) (**Supplemental Figure 6A, D**). We generated a mutant, *smfD*_Δpatch_, that lacks these two residues and the ten amino acid sequence between them to potentially generate a SmfD variant that would minimally alter protein structure, especially the Smf-1 binding region, but disrupt aggregation. Transmission electron microscopy (TEM) revealed that compared to the WT, both Δ*smf-1* and Δ*smfD* exhibited a smoother outer membrane devoid of fimbriae (**Supplemental Figure 7**). It is likely that the absence of the distal SmfD adhesin leads to potential destabilization of the Smf-1 fimbriae, as has been seen in other fimbrial systems (*67*). In contrast, *smfD*_Δpatch_ cells exhibited numerous fimbriae that appeared longer and more tangled than WT fimbriae, indicating putative aberrant fimbrial formation in the *smfD*_Δpatch_ mutant.

To test whether these fimbriae were functional, we first assayed the production of biofilms, as Smf-1 is known to be required for biofilm formation (*55*). Unlike Δ*smf-1* and Δ*smfD*, which were both biofilm-negative, the *smfD*_Δpatch_ mutant had only a partial defect in biofilm formation (**Supplemental Figure 6E**), indicating that the *smfD*_Δpatch_ fimbriae retain some function. Further, we found that the *smfD*_Δpatch_ mutant formed aggregates both in the absence and presence of *Pa* secreted molecules, indicating that the aberrant fimbriae in this mutant are ‘sticky’, and cause cells to adhere to each other (**Supplemental Figure 6F, G**). However, we found that in a CFU enumeration assay, the *smfD*_Δpatch_ mutant was functionally indistinguishable from the Δ*smf-1* and Δ*smfD* mutants, as it showed similar CFUs to those mutants with and without exposure to *Pa* supernatant, indicating that the aggregates formed by this mutant are likely less adherent, and unable to withstand the minimal agitation they are subject to in this assay (**Supplemental Figure 6H**). Cumulatively, these data suggest that the SmfD adhesin is an important element in facilitating aggregation of *Sm* in response to *Pa* secreted factors.

### *Pa* siderophores, rhamnolipids, and quorum sensing indirectly mediate *Sm* aggregation

*Pa* has an extensive and well-described secretome and many of these molecules have been shown to exert toxic effects or alter behavior and physiology of competing bacteria (*22*, *68–70*). To test if one of these known secreted factors was responsible for *Sm* aggregation, we derived supernatant from *Pa* mutants deficient in the production of the surfactant rhamnolipids (Δ*rhlA*), iron-chelating siderophores pyoverdine and pyochelin (Δ*pvdJ* Δ*pchE*), alkyl-quinolones (Δ*pqsA*), redox-active phenazines (Δ*phz1/2*), the LasA protease (Δ*lasA*), and hydrogen cyanide (Δ*hcnABC*). We found that while supernatants from the other mutants were similar to that of WT *Pa,* supernatants from the siderophore (Δ*pvdJ* Δ*pchE*), rhamnolipid (Δ*rhlA*), and alkylquinolone (Δ*pqsA*) mutants induced reduced *Sm* aggregation, as measured by higher *Sm* CFUs, especially at the 4- and 8-hour time points (**Supplemental Figure 8A**).

We first examined the role of the *Pa* siderophores and iron signaling in inducing *Sm* aggregation. Neither elevated nor reduced iron concentrations were found to induce *Sm* aggregation as assessed through exposure to excess ferrous/ferric iron or diverse iron chelators, respectively. (**Supplemental Figure 8B**). Further, addition of pyoverdine directly to *Sm* or to supernatant from the Δ*pvdJ* Δ*pchE* mutant also did not induce *Sm* aggregation, but secreted products from the Δ*pvdJ* Δ*pchE* mutant grown with exogenous pyoverdine restored the induction of *Sm* aggregation (**Supplemental Figure 8C**). These findings indicate that siderophore signaling plays an important but indirect role in mediating *Pa*’s ability to induce aggregation in *Sm*.

We saw similar results with the biosurfactant rhamnolipid, and the *N*-butyryl-homoserine lactone (C4-HSL) and *N*-3-oxo-dodecanoyl-L-homoserine lactone (3-oxo-C12-HSL) quorum sensing molecules. Supernatants derived from mutants in the biosynthesis pathways for each of these molecules, Δ*rhlA* for rhamnolipids, Δ*rhlI* for C4-HSL, and Δ*lasI* for 3-oxo-C12-HSL, were deficient in *Sm* aggregation induction (**Supplementary** Figures 8D-F). However, addition of the respective exogenous molecules to *Sm* was not sufficient to restore *Sm* aggregation. Supernatant from the *Pa* mutants grown with the respective molecules induced aggregation, indicating an indirect effect of these molecules on the production of the aggregation inducing factor by *Pa*.

### PQS induces *Sm* aggregation

Having identified several indirect modulators required for *Pa* to induce *Sm* aggregation, we next sought to identify *Pa* determinants that directly induced *Sm* aggregation. We fractionated concentrated *Pa* supernatant and assessed each fraction for aggregation induction in *Sm*. We found that of 19 fractions, two—fractions 11 and 12— induced robust *Sm* aggregation (**Figure 4A, Supplementary Figures 9A, B**). Mass spectrometry analysis of the most active fractions 11 and 12 in comparison to adjacent fractions 10 and 13, respectively, revealed a variety of enriched molecules within the active fractions (**Supp. Table 3**). In particular, 2-heptyl-3-hydroxy-4-quinolone (PQS) was enriched 11-fold in fraction 11 compared to neighboring fraction 10 and 6-fold in fraction 12 compared to fraction 13 (**Figure 4B**), consistent with the hypothesis that PQS within *Pa* supernatant may induce *Sm* aggregation.

**Figure 4.**
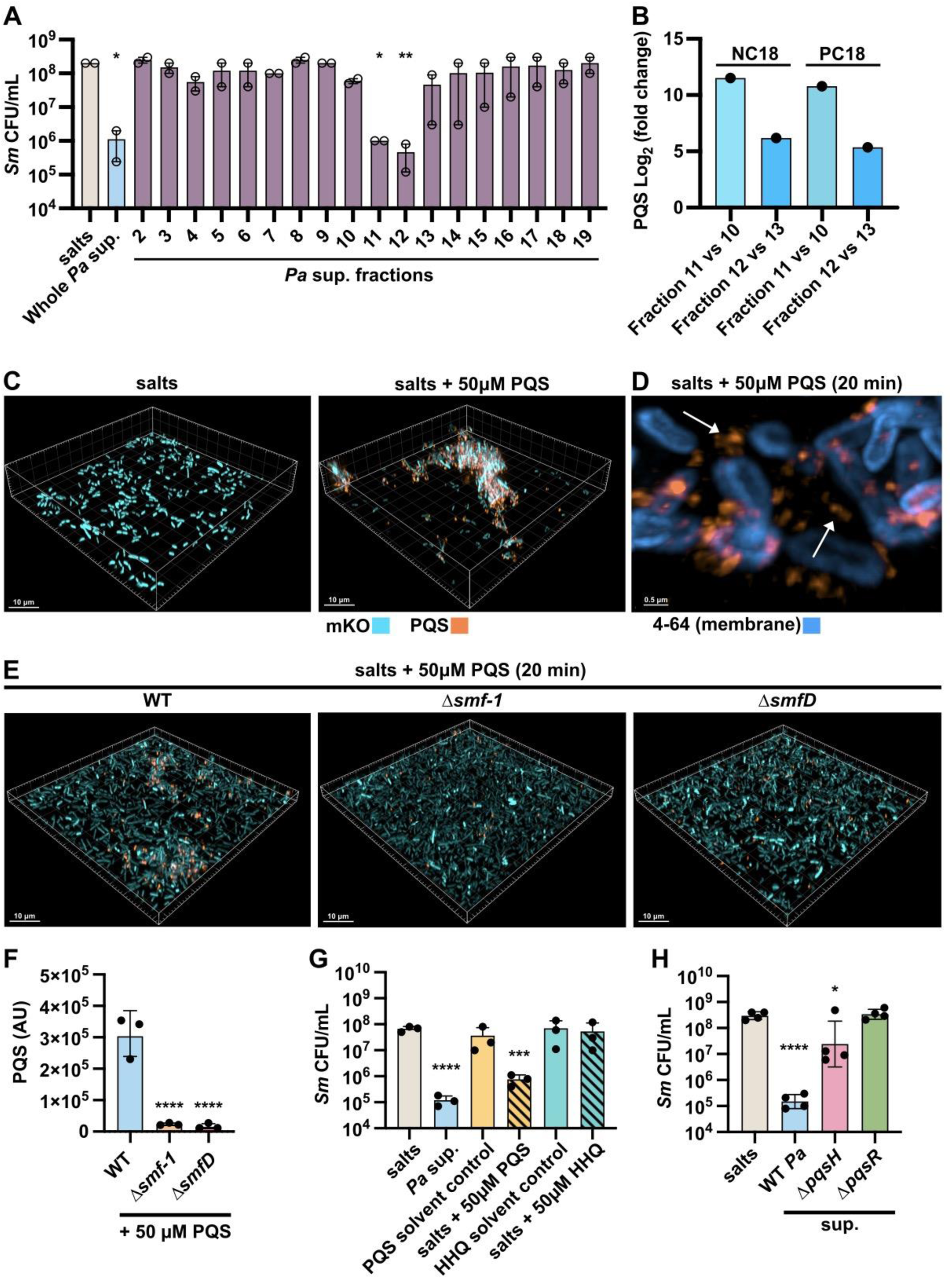
PQS induces *Sm* aggregation. **(A)** CFU enumeration of *Sm* exposed to salts, concentrated whole *Pa* cell-free supernatant and fractions from concentrated *Pa* supernatant for 2 h. **(B)** Log2 (fold-change) of PQS enrichment in fractions 11 versus 10 and fraction 12 versus 13 as measured via QToF-MS/ UHPLC in C18 columns, under both positive (PC18) and negative (NC18) electrospray conditions. **(C)** Microscopy of mKO tagged *Sm* cells exposed to salts control and salts supplemented with 50 μM exogenous PQS for 2 h. **(D)** Microscopy of *Sm* stained with membrane dye FM™ 4-64 (blue) and imaged after 20 min exposure to salts with 50 μM exogenous PQS supplementation. White arrows indicate extracellular PQS signal. **(E)** Microscopy of mKO tagged *Sm* WT, Δ*smf-1*, and Δ*smfD* after 20 min exposure to a salts control or salts supplemented with 50 μM exogenous PQS. **(F)** Quantification of PQS fluorescence intensity normalized to total biovolume in *Sm* WT, Δ*smf-1*, and Δ*smfD* after 20 min exposure to 50 μM exogenous PQS. **(G)** CFU enumeration of *Sm* exposed to salts, solvent controls, *Pa* supernatant, 50 μM exogenous PQS, and 50 μM exogenous HHQ for 2 h. **(H)** CFU enumeration of *Sm* exposed to salts, and supernatant derived from *Pa* WT, Δ*pqsH*, or Δ*pqsR* for 2 h. **(A, F-H)** Data shown are for **(A)** two or **(F-H)** three biological replicates and represented as the mean ± SD. Significance is shown for comparison to **(A, G, H)** the respective salts control and **(F)** the respective WT control as tested by a one-way ANOVA on **(A, G, H)** log-transformed or **(F)** measured values followed by the Tukey’s test for multiple comparisons (*, *p* < 0.05; **, *p* < 0.01; ***, *p* < 0.001; ****, *p* < 0.0001). **(C-E)** Representative images of three biological replicates are shown.

We found that the addition of 50 μM exogenous PQS led to robust *Sm* aggregation. In addition, we could detect PQS due to its intrinsic fluorescence (*71*, *72*), and observed that PQS localized within and around *Sm* aggregates (**Figure 4C**). Further, staining with the membrane dye FM™ 4-64 in conjunction with single-cell imaging showed PQS to be localized external to the *Sm* membrane (**Figure 4D**). PQS localization to *Sm* aggregates was significantly reduced in both the Δ*smf-1* and Δ*smfD* mutants compared to WT, indicating a role of the Smf-1 fimbriae in the PQS association (**Figure 4E, F**).

Production of PQS requires the transcriptional regulator PqsR (MvfR), the *pqsABCDE* biosynthetic operon, which produces alkyl-quinolone intermediates, and PqsH, which converts the final intermediate, HHQ, into PQS (*73*). We found that while exogenous PQS induced *Sm* aggregation as also measured by the characteristic decrease in *Sm* CFUs, addition of exogenous HHQ had no impact (**Figure 4G**), indicating that *Sm* aggregation was a specific response to PQS. The reduction in *Sm* CFUs observed after PQS exposure was specifically due to aggregation, and not cell death, as dispersal of PQS-induced aggregates exhibited similar CFU values as the dispersed salts control (**Supplemental Figure 9C**). Finally, supernatant derived from Δ*pqsH* and Δ*pqsR* mutants, which are both unable to synthesize PQS, failed to induce *Sm* aggregation (**Figure 4H**). In sum, we find that the *Pa* quorum-sensing molecule PQS is necessary and sufficient to induce *Sm* aggregation, and associates with *Sm* aggregates in an Smf-1 fimbriae-dependent manner.

### *Sm* Smf-1 mediated aggregation is protective against *Pa* in co-culture

We next assessed the potential consequences of this interspecies interaction. *Pa* is known to exert contact-dependent and independent antagonism against several species (*3*, *74–76*). We hypothesized that the interspecies interaction between *Pa* and *Sm* described here is part of a defensive multicellular behavioral mechanism through which *Sm* is afforded protection from *Pa*-mediated antagonism. To test this, we conducted co-culture assays on an agar surface, an environment where *Pa* has been shown to readily compete through invasion and microcolony disruption (*77*). First, we separately overlaid *Sm* WT, Δ*smf-1*, or Δ*smfD* with *Pa* WT and after 4 h of co-culture, observed that *Sm* WT formed tight multicellular structures, while *Pa* primarily localized to the periphery (**Figure 5A**). In contrast, co-cultures of *Sm* Δ*smf-1* or Δ*smfD* with *Pa* both exhibited intermixing of single cells from both species and the absence of microcolony or aggregate structures. After 16 h of co-culture, *Sm* WT exhibited bacterial aggregates, while the Δ*smf-1* and Δ*smfD* co-cultures remained in single-cell form and were overgrown by *Pa* (**Figure 5B**). In contrast, *Sm* WT co-cultured with *Pa* Δ*pqsR* exhibited significantly reduced aggregation (**Figure 5C**). *Sm* WT showed significantly higher survival compared to either the Δ*smf-1* or Δ*smfD* mutants after 16 h of co-culture with *Pa* WT, but not in monoculture controls (**Figure 5D, E**). Further, while we observed higher CFU counts of all *Sm* backgrounds when grown in co-culture with *Pa* Δ*pqsR*, consistent with a competition defect, these counts were still lower than the monoculture controls, but *Sm* WT did not have a survival advantage compared to the Δ*smf-1* or Δ*smfD* mutants (**Figure 5F**). These findings demonstrate that *Sm* aggregation is a protective response in co-culture with *Pa*.

**Figure 5.**
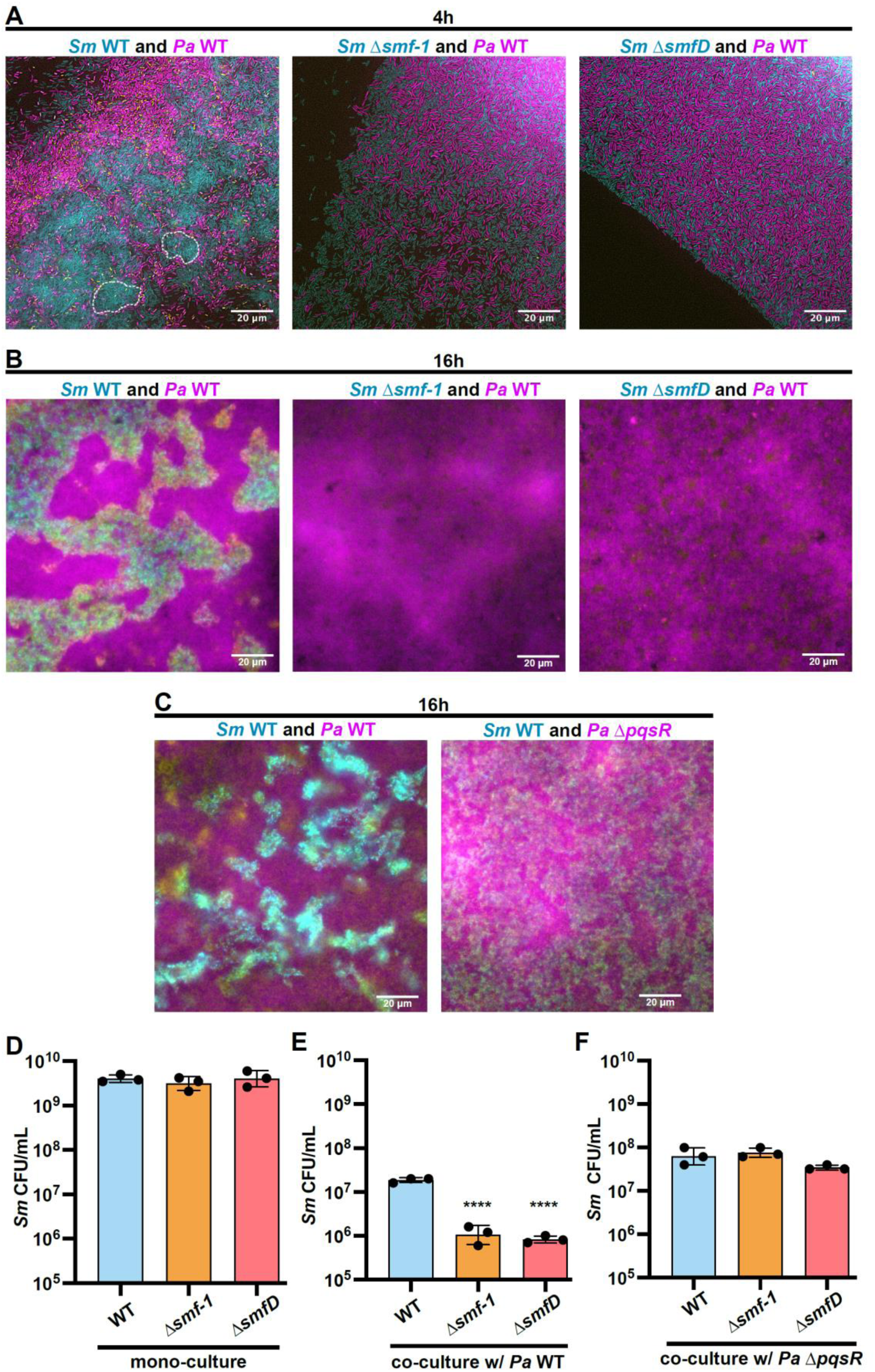
*Sm* aggregation provides a survival advantage from *Pa* competition. **(A, B)** GFP tagged *Sm* WT, Δ*smf-1*, and Δ*smfD* were grown on agar for 4 h and subsequently overlaid with mKO tagged *Pa* WT and agar cutouts were imaged after an additional co-culture growth for **(A)** 4h and **(B)**16 h co-culture growth. Examples of early aggregates of *Sm* WT are marked by white dotted lines. **(C)** Similar 16 h co-culture of GFP tagged *Sm* WT with mKO tagged *Pa* WT or Δ*pqsR*. **(A-C)** Representative images of three biological replicates are shown. **(D-F)** CFU enumeration following mechanical dispersion of agar cultures of Sm WT, Δ*smf-1*, and Δ*smfD* **(D)** in mono-culture, or in co-culture with *Pa* **(E)** WT, or **(F)** Δ*pqsR*. Data shown are the mean ± SD for three biological replicates. Significance is shown for comparison to the respective WT control as tested by a one-way ANOVA on log-transformed values followed by the Tukey’s test for multiple comparisons (****, *p* < 0.0001).

Next, to assess the contribution of the Smf-1 fimbriae specifically to invasion resistance, we used a macroscopic colony invasion model where we inoculated *Pa* cultures adjacent to *Sm*. After 24 h, we observed that *Pa* swarmed and grew peripherally around the *Sm* WT microcolony, which maintained its integrity (**Figure 6A-C**). In contrast, *Pa* readily breached the periphery of Δ*smf-1* and Δ*smfD* microcolonies and subsequently invaded their core. This was reflected in *Sm* fitness, where *Sm* WT had significantly higher survival compared to the Δ*smf-1* and Δ*smfD* mutants (**Figure 6D**). These data show that the Smf-1 fimbriae mediate aggregation and resistance to colony invasion, and are critical determinants for *Sm* survival in co-culture with *Pa*.

**Figure 6.**
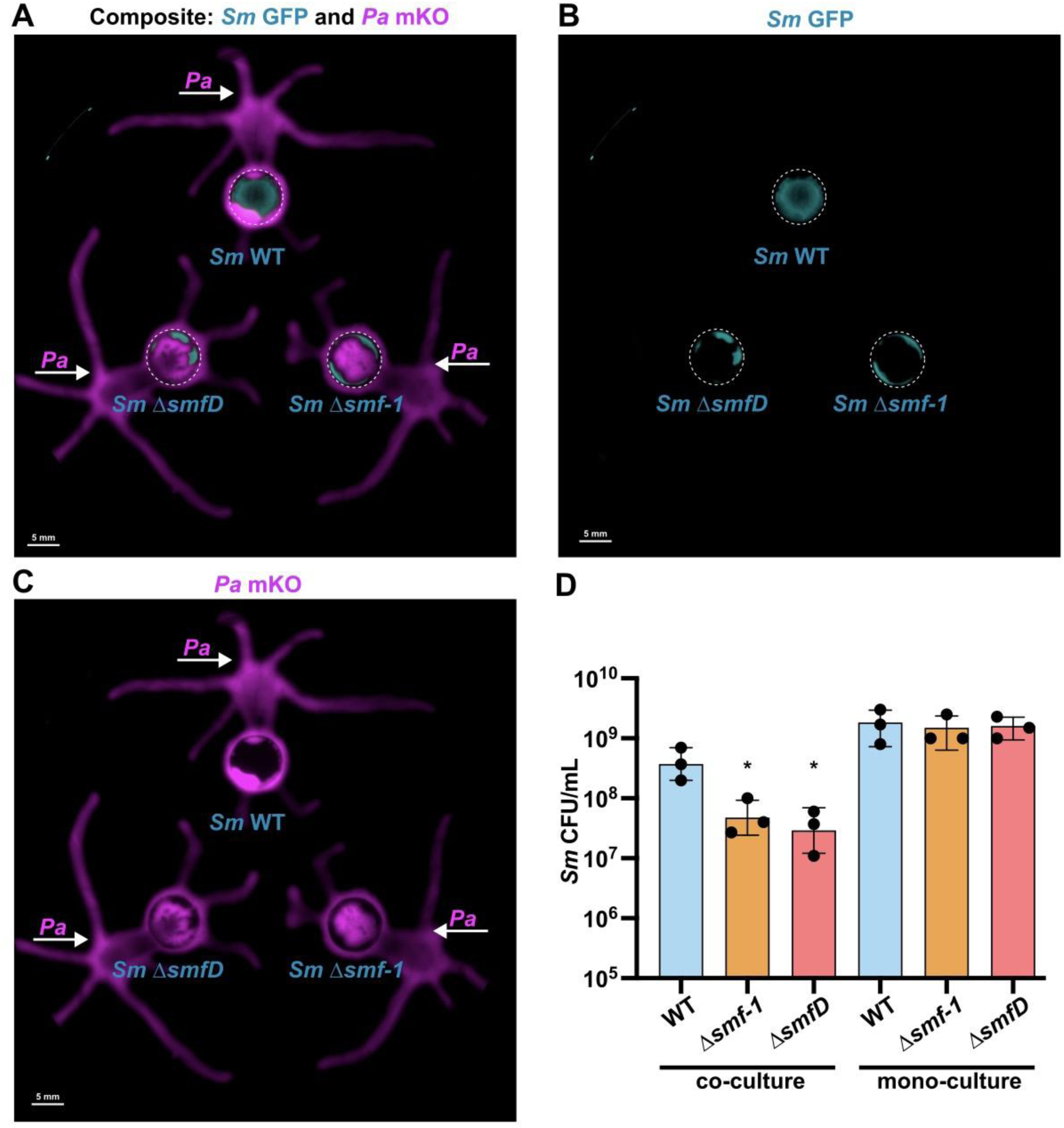
Smf-1 fimbriae mediate *Sm* resistance to *Pa* colony invasion. Imaging of GFP tagged *Sm* WT, Δ*smf-1*, and Δ*smfD* spotted on an agar plate for 4 h prior to adjacent inoculation of mKO tagged PA14 WT and growth for an additional 24 h shown for **(A)** composite signals, **(B)** GFP channel, and **(C)** mKO. Agar inoculation of *Sm* GFP strains and PA14 mKO indicated via dashed circles and white arrows, respectively. **(D)** Enumeration of co-culture of *Sm* WT, Δ*smf-1*, and Δ*smfD* with *Pa* WT, and monoculture controls after removal and mechanical dispersion. Data shown are the mean ± SD for three biological replicates. Significance is shown for comparison to the respective WT control as tested by a one-way ANOVA on log-transformed values followed by the Tukey’s test for multiple comparisons (*, *p* < 0.05). **(A-C)** Representative images of three biological replicates are shown.

### Interspecies metabolite-induced aggregation is observed in additional bacterial species

To assess whether *Pa* supernatant-induced aggregation was specific to *Sm* or also observed in additional bacterial species, we challenged a number of Gram-negative bacteria with *Pa* supernatant and evaluated aggregation macroscopically and microscopically. Out of 7 isolates from 4 different species, we found evidence of *Pa* supernatant-induced aggregation in the single isolate of *Salmonella enterica* and all three isolates of *Achromobacter xylosoxidans*, but not in isolates of *E. coli* and *Burkholderia cenocepacia* (**Figure 7**), indicating that aggregation is a more general, but not ubiquitous bacterial response to *Pa* secreted factors.

**Figure 7.**
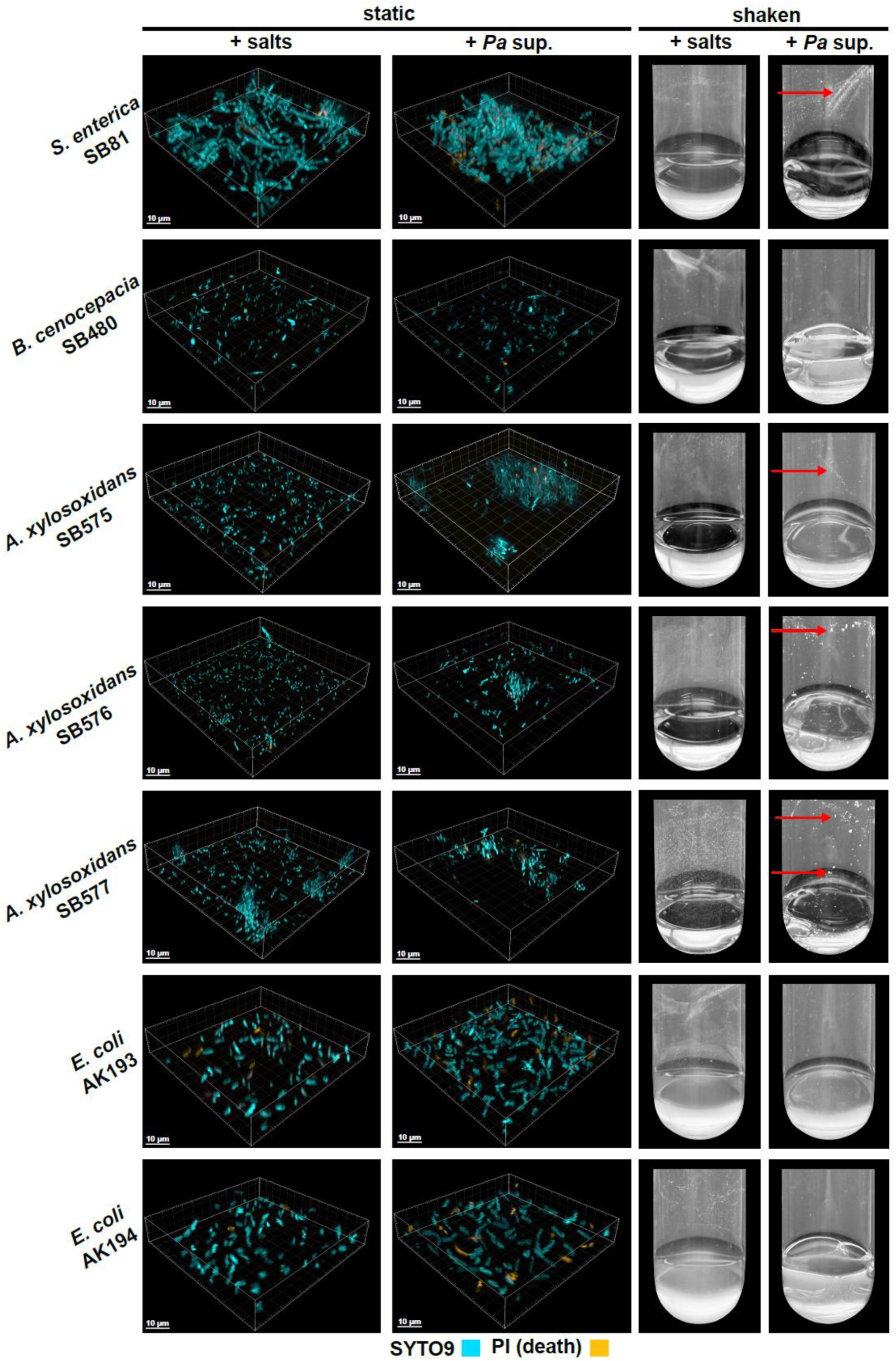
Interspecies metabolite-induced aggregation is observed in additional bacterial species. Early-log *Salmonella enterica, Burkholderia cenocepacia, Achromobacter xylosoxidans,* and *Escherichia coli* cells were challenged with 50% (v/v) *Pa* cell-free supernatant. **(A)** Microscopy images after 2 h incubation followed by SYTO9 and propidium iodide (PI)-staining. **(B)** Images of shaken tubes after 30 minutes incubation. Macroscopic aggregates are indicated with red arrows.

## DISCUSSION

We provide evidence of a novel interspecies interaction between the co-infecting pathogens *S. maltophilia* and *P. aeruginosa*, finding that *S. maltophilia* utilizes Smf-1 fimbriae to mediate defensive multicellularity in response to *P. aeruginosa*-secreted factors (**Figure 8**). Similar aggregation by *S. enterica* and *A. xylosoxidans* in response to *Pa* secreted factors suggest that this interaction may be a more general phenomenon.

**Figure 8.**
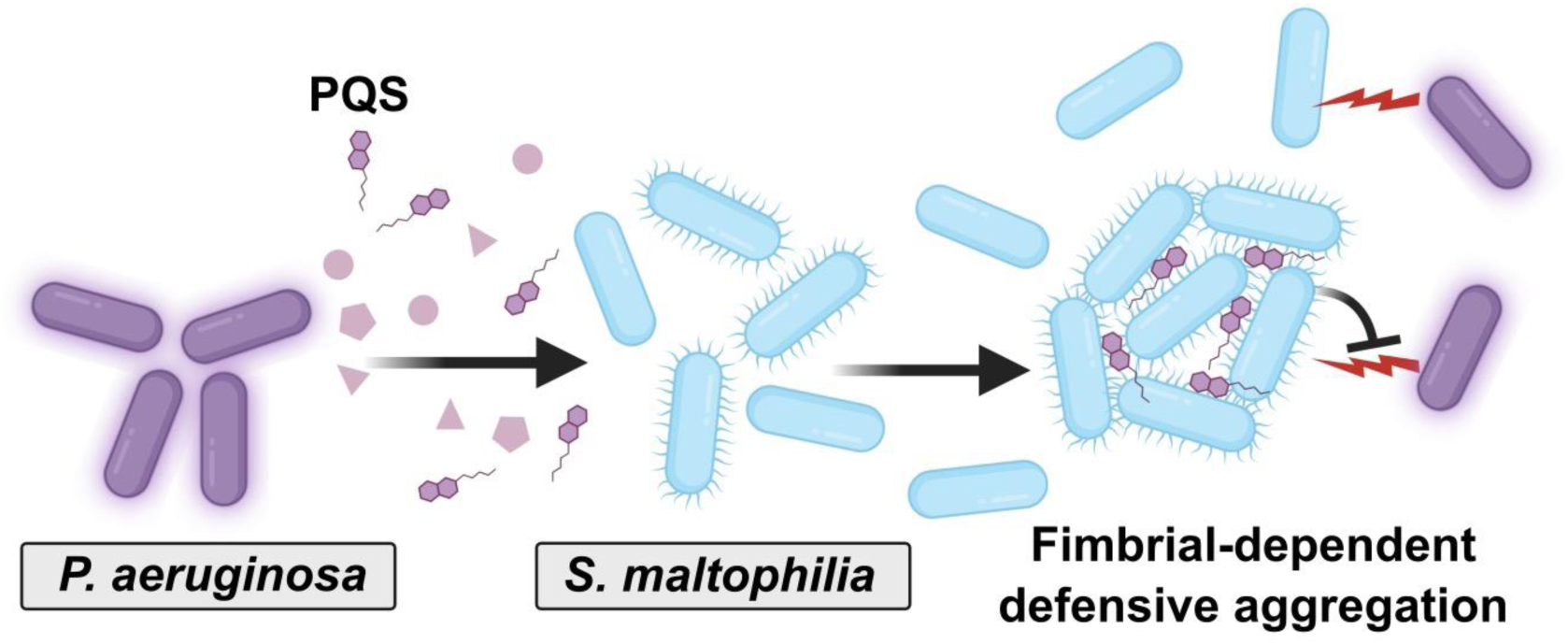
A model for interspecies metabolite-induced aggregation. *S. maltophilia* (blue rods) aggregate in response to *P. aeruginosa*-produced PQS (purple hexagons). *S. maltophilia* requires the Smf-1 fimbriae to aggregate, a phenotype which enhances protection from *Pa*-mediated antagonism.

*Sm* aggregation required the presence of Smf-1 fimbriae on both partnering cells, consistent with bridging aggregation that occurs when surface-attached and free-floating molecules or complexes such as fimbriae or polysaccharides, act as bridges between different cells (*78–80*). Aggregation and biofilm formation utilize many shared determinants such as the presence of fimbriae and pili to mediate initial attachment and the production of extracellular polysaccharide (EPS) to collectively enmesh bacteria and provide protection from exogenous threats such as antibiotics, predation, and phagocytosis (*81–85*). While we do not exclude a contribution of EPS to *Sm* aggregation, the short timeframe (<15 minutes) required for the induction of this phenotype in response to *Pa* secreted factors, and the lack of co-aggregation between WT and Δ*smf-1* cells, suggests that the Smf-1 fimbriae alone are sufficient for this behavior.

We found exogenously added PQS induced *Sm* aggregation with microscopy consistent with extracellular localization of this quorum sensing molecule within *Sm* aggregates. PQS is a critical component of *P. aeruginosa*’s quorum sensing systems, and its production and signaling are required for *P. aeruginosa* biofilm morphology, the release of extracellular DNA (eDNA), outer membrane vesicle biogenesis, and the secretion of virulence factors (*86–89*). *Pa* supernatant from the LasI mutant was unable to induce *Sm* aggregation (**Supp.** Fig. 8F), in agreement with previous reports finding PQS production to be positively regulated by Las quorum sensing (*90*). However, supernatant from the *Pa* Δ*pqsA* mutant that is completely deficient for the production of all PQS family molecules only partially abrogated *Sm* induction (**Supp.** Fig. 8A), suggesting that additional *Pa* secreted factors may also be involved in this phenotype. Rhamnolipids, siderophores, and the Rhl quorum sensing system may indirectly alter the production of either PQS or these additional factors, leading to the aggregation induction defect seen in the respective mutants (**Supp.** Fig. 8).

In addition to its role as a quorum sensing and signaling molecule in *Pa*, PQS has been shown to alter interspecies behavior and physiology, partially due to its capacity to chelate iron (*91*). PQS inhibits motility in a number of gram-negative and positive bacterial species alike, while other studies have demonstrated PQS-mediated inhibition of *Streptococcus mutans* biofilm formation, enhanced membrane vesicle production in both *E. coli* and *Mycobacterium abscessus*, and lysis and growth inhibition of bacterial soil isolates (*92–97*). PQS has been shown to directly bind elements of the *Pa* outer membrane such as the efflux pump protein MexG, O-antigen, LPS, and components of the Type 6 secretion system, and given the presence of PQS within *Sm* aggregates, we hypothesize that PQS similarly bind *Sm’s* Smf-1 fimbriae (*72*, *98*). Further work will explore this putative binding, as well as any signaling mediated by PQS.

We propose that *Sm* aggregation is a defensive multicellular behavioral response to PQS, which is recognized as an exogenous foreign danger signal. Danger sensing pathways in bacteria are thought to sense self or foreign cues that indicate a potential threat (*17*), and mount a protective or antagonistic response. Self-danger cues include peptidoglycan and norspermidine released through kin cell lysis, both of which lead to increased biofilm production (*15*, *16*).

Induction of defensive bacterial behavior upon sensing of foreign cues as danger signals remains poorly characterized. Notably, danger sensing is distinct from competition sensing where bacteria sense stresses potentially derived from competing microbes and respond antagonistically (*19*). While competition sensing typically is a response to cellular damage or nutrient limitation, danger sensing responds to imminent threats (*17*). Foreign quorum-sensing molecules can lead to non-specific fitness alterations or an increased competitiveness index (*99*, *100*), but specific protective responses to interspecies quorum sensing are not well defined. A quorum sensing molecule is an especially informative cue as it not only signals the presence of a competitor but also indicates high cell density of the competitor and potentially the presence of associated antimicrobial virulence factors. The response of *Sm* to PQS is also very specific as we did not see a similar response to HHQ, the immediate precursor to PQS and a structurally very similar molecule.

Danger signals are perceived as imminent threats, and do not directly induce stress or lead to loss of viability. Our work shows that PQS does not cause cell death in *Sm*, and stresses like hypoxia, and ATP or iron depletion do not lead to aggregation, supporting the idea that PQS is sensed as a danger signal, and that aggregation is not a stress response. However, further work will investigate whether PQS leads to other stresses, or whether additional physicochemical stresses, like reactive oxygen species, or osmotic or pH changes, can induce aggregation.

The mechanism by which PQS is sensed by *Sm* remains under investigation. Non-exclusive possibilities include physical interactions with *Sm* Smf-1 fimbriae, or induction of a signaling cascade, which would explain the lack of robust aggregation in the absence of protein translation. Future studies will focus on the signaling mechanisms and active processes required for the sensing of PQS by *Sm* and its subsequent aggregation response, the regulation of Smf-1 fimbriae and interactions with PQS, and whether these processes are conserved in interspecies aggregation by additional species.

Smf-1-dependent *Sm* aggregation in response to *Pa* PQS expands our understanding of the role of quorum sensing molecules in interspecies interactions and the contribution of extracellular appendages in effecting multicellular defensive behavior in response to foreign danger cues. Our work demonstrates that interspecies interactions can thus modulate novel bacterial behaviors, sensing, and protective or competitive responses, which likely significantly alter microbial fitness and community dynamics in polymicrobial settings.

## MATERIALS AND METHODS

### Bacterial strains and growth conditions

Bacterial strains used in this study are listed in **Supp. Table 4**. *P. aeruginosa* UCBPP-PA14 (*101*) and *S. maltophilia* K279a (*102*) and their derivatives were grown in a modified M63 medium (*103*) made of 13.6 g·L^-1^ KH_2_PO_4_, 2 g·L^-1^ (NH_4_)_2_SO_4_, 0.4 µM ferric citrate, 1 mM MgSO_4_; pH adjusted to 7.0 with KOH and supplemented with 0.3% glucose, 1× ACGU solution (Teknova), 1× supplement EZ (Teknova), 0.1 ng·L^-1^ biotin, and 2 ng·L^-1^ nicotinamide, at 37°C, with or without shaking at 300 rpm when specified. For cloning and strain construction, strains were cultured in Luria Bertani (Miller) broth or on LB agar plates with 15 g·L^-1^ agar, supplemented with 50 µg·mL^-1^ gentamicin, 30 µg·mL^-1^ chloramphenicol, or 6.25 or 25 µg·mL^-1^ irgasan for *S. maltophilia* or *P. aeruginosa*, respectively.

### Media Supplementation

The following chemicals were used to test their effect on *S. maltophilia* aggregation: 0.1% gastric porcine mucin (Sigma-Aldrich), 50 μM 2-heptyl-3-hydroxy-4-quinolone (PQS) (Santa Cruz Biotechnology), 50 μM 2-heptyl-4-hydroxyquinoline (HHQ) (Sigma-Aldrich), 50 μM pyoverdines derived from *Pseudomonas fluorescens* (Sigma-Aldrich), µg·mL^-1^ rhamnolipids (Sigma-Aldrich), 1 μM *N*-butyryl-homoserine lactone (C4-HSL) (Cayman Chemical Company), 1 μM *N*-3-oxo-dodecanoyl-*L*-homoserine lactone (3-oxo-C12-HSL) (Sigma-Aldrich), 100 μM ferric citrate (Sigma-Aldrich), 100 μM ferrous chloride (Sigma-Aldrich), 100 μM ferrozine (Sigma-Aldrich), 100 μM conalbumin (Sigma-Aldrich), and 100 μM 2,2’-bipyridyl (Sigma-Aldrich).

### Preparation of cell-free supernatant

Overnight cultures of *P. aeruginosa* (or *S. aureus* JE2 and *S. maltophilia* K279a for the data shown in **Figure 1A**) were diluted to an OD_600_ of 0.05 in fresh media and grown for 16 h in flasks before harvesting at 4,000 rpm for 30 minutes. The supernatant was sterile filtered with a Steriflip with a 0.22 μm polyethersulfone filter (MilliporeSigma). *For P. aeruginosa* strains grown with specific chemicals, mutant strains were grown overnight in media individually supplemented with 50 μM pyoverdines, 100 µg·mL µg·mL^-1^ rhamnolipids, 1 μM C4-HSL, or 1 μM 3-oxo-C12-HSL. These overnight cultures were diluted to an OD_600_ of 0.05 in fresh media containing the respective supplement and grown in a flask for 16 h prior to supernatant filtration as above. Supernatants were stored at - 30°C until use.

### Quantification of *S. maltophilia* CFUs upon exposure to *P. aeruginosa* cell-free supernatant

Overnight cultures of *S. maltophilia* were diluted to an OD_600_ of 0.05 in fresh media and grown in a flask to an OD_600_ of 0.15. At this time, 500 μL of culture was added to 500 μL *P. aeruginosa* cell-free supernatant (or derivatives of supernatant or secreted products) or media salts as a control in 14 ml culture tubes. After incubation at different time points at 37°C accompanied by shaking at 300 rpm, serial dilutions of the *S. maltophilia* culture were spotted on LB agar plates and enumerated the next day. To mechanically disperse bacterial aggregates, bacteria in tubes were vortexed for 10 seconds to remove adherent aggregates and then homogenized with a 1 mL syringe fitted with a 27G 0.5-inch needle prior to enumeration.

### Microscopy

Overnight cultures of *S. maltophilia* were diluted to an OD_600_ of 0.05 in fresh media and grown in a flask to an OD_600_ of 0.15. Then, 50 μL of culture was added to 50 μL *P. aeruginosa* supernatant or M63 salts medium control and incubated for 2 h statically at 37°C. Upon addition to the 96-well glass bottom plate (Cellvis), bacteria were stained with 10 µg·mL^-1^ propidium iodide (PI) to assess cell death, 1 µg·mL^-1^ FM4-64 to visualize cell membranes, and 5 μM SYTO9 as needed. Epifluorescence microscopy of aggregates of *S. maltophilia* and additional species were conducted at room temperature with a DeltaVision microscope (Applied Precision/GE Healthcare), Olympus UPlanSApo 100x/1.40 oil objective, and an EDGE sCMOS 4.2 camera using a 12 μm Z-stack. For the data shown in **Figure 4D-E**, a 4 μm Z-stack was used. Micrographs were deconvolved with SoftWorx software (GE Healthcare), and Imaris 10.1 was used to render, manually segment, and quantify object biovolumes using the surfaces function and employing a 5 μm local contrast background subtraction.

### Macroscopic tube aggregation assays

Overnight cultures of all species were diluted to an OD_600_ of 0.05, grown in fresh media to an OD_600_ of 0.15, exposed to 50% (v/v) M63 medium salts control or *P. aeruginosa* supernatant, and shaken at 37°C for 15 minutes (*S. maltophilia* data shown in **Figure 1B**) or 60 minutes (other species data shown in **Figure 7**). At the designated timepoint, tubes were removed from 37°C and imaged. To assay for aggregation inhibition, *S. maltophilia* was grown in M63 medium as previously described. Prior to exposure to *P. aeruginosa* supernatant, *S. maltophilia* was inoculated into 96-well glass bottom plates and challenged for 20 minutes by either static placement in an anaerobic chamber, 20 μM Carbonyl Cyanide *m*-Chlorophenylhydrazone (CCCP), or 20 µg·mL^-1^ Chloramphenicol. After the challenge, *S. maltophilia* was exposed to 50% (v/v) *P. aeruginosa* supernatant, stained with PI, and experimental conditions were maintained for an additional 2 h, prior to epifluorescence microscopy.

### Quantification of aggregation and live/dead ratios

Biovolumes of all Imaris-segmented objects were quantified from a minimum of three biological replicates per experiment. Aggregates observed in epifluorescence microscopy-derived images were defined as any object with a volume greater than 20 μm^3^ and calculated as a percentage of the total biovolume of bacteria imaged. Cell death percentage was calculated through biovolume quantification of PI-stained cells as a percentage of the biovolume of all cells, using either constitutive fluorescent GFP tags or staining with 5 μM SYTO9 as indicated. Biovolumes of all Imaris-segmented objects were quantified from a minimum of three biological replicates per experiment.

### Transmission Electron Microscopy

Overnight cultures of *S. maltophilia* were diluted to an OD_600_ of 0.05, grown in fresh media to an OD_600_ of 0.15, back-diluted 1:10, and fixed in a 4% (v/v) solution of PBS-paraformaldehyde. Samples were negatively stained with 0.5% phosphotungstic acid and visualized with an HT7800 electron microscope (Hitachi) under 8,000-30,000X magnification operating at an accelerating voltage of 8 kV under high vacuum.

### Experimental evolution of *S. maltophilia* in *P. aeruginosa* supernatant

Overnight cultures of *S. maltophilia* were diluted to an OD_600_ of 0.03 in 10 mL flasks and grown to an OD_600_ of 0.15. At this time, 500 μL of culture was added to culture tubes with 500 μL *P. aeruginosa* supernatant. After incubating the culture in WT *P. aeruginosa* supernatant for 16 h which allows for recovery of cell numbers, the resulting *S. maltophilia* cultures were used to inoculate a new 10 mL flask at OD_600_ of 0.03 to start the process over again. Two independent populations were passaged, and isolates were obtained by streaking out the populations and picking single colonies. Three isolates each were obtained from passages 5 and 10 for population 1, and passage 5 from population 2.

### Whole genome sequencing and analysis

The nine evolved isolates of *S. maltophilia* described above, as well as the WT K279a strain were analyzed by whole genome sequencing (WGS). Bacterial cultures were grown overnight in LB broth and 1 mL was harvested for genomic DNA purification using the DNeasy Kit (Qiagen) according to the manufacturer’s instructions. WGS was performed at the NCI Center for Cancer Research Genomics Core, using Illumina sequencing. The sequencing files were processed with Cutadapt (*104*) and Trimmomatic (*105*). The mutations in the evolved isolates were identified using breseq (*106*) and are listed in **Supp. Table 1**.

### Bioinformatics and modeling

DELTA-BLAST of *S. maltophilia* SmfD’s receptor binding domain (41-187) was used to identify putative homologs, *Klebsiella pneumoniae* MrkD and *Acinetobacter baumannii* Abp1D (*107*) and CLUSTALW was used to align sequences (*108*) . The structural models were generated by submitting the protein sequences to the AlphaFold Server (*109*) and default settings were applied. Then ChimeraX (*110*) was used to examine the top 5 models and the ones most consistent with our experimental data were depicted.

### RNA preparation and sequencing

Overnight cultures of *S. maltophilia* were diluted to an OD_600_ of 0.05 and grown to an OD_600_ of 0.15, prior to exposure to 50% (v/v) M63 salts medium control or *P. aeruginosa* supernatant, for 30 minutes. 2 mL of these cultures was mixed with 4 mL of RNAprotect Bacteria Reagent (Qiagen) and incubated for 5 min at room temperature followed by centrifugation at 4,000 rpm for 10 min. Supernatants were decanted and pellets stored at −80°C. RNA was extracted using the Total RNA Purification Plus Kit (Norgen) per the manufacturer’s instructions for Gram-negative bacteria and DNase I treated to remove genomic DNA using the TURBO DNA-free Kit (Invitrogen). A PCR was used to verify the absence of contaminating DNA. Integrity of the RNA was confirmed by running an aliquot of the RNA sample on an agarose gel to observe rRNA bands. Finally, rRNA was removed using the *S. maltophilia* riboPOOLs rRNA Depletion Kit (siTOOLs Biotech) followed by library preparation with the NEBNext Ultra II Directional RNA Library Prep Kit for Illumina (New England Biolabs). Sequencing was performed at the Center for Cancer Research (CCR) Genomics Core Facility. Two independent biological replicates were collected.

### RNA-seq analysis

Adapter sequences were removed from reads in the FASTQ files using Cutadapt (*104*) and reads were further processed with Trimmomatic (*105*). We aligned the reads to the *S. maltophilia* K279a genome (NC_010943.1) using Rockhopper (*111–113*) . After removal of reads mapping to the rRNA genes, raw read counts were analyzed with DESeq2 (*114*) to obtain the fold-change in gene expression between the salts and *P. aeruginosa* cell-free supernatant conditions, and statistical significance of the comparisons. Pathway analysis was performed using FUNAGE-Pro (*115*) on lists of significantly upregulated or downregulated genes, using a cut-off of fold-change in gene expression > 2 or < 0.5 respectively, and an adjusted *p-*value < 0.05.

### Strain construction

Homologous downstream and upstream arms of the respective WT or mutant gene alleles were amplified using the primers listed in **Supp. Table 5**. For *S. maltophilia*, amplified fragments were cloned into pDONRpEX18Gm (*116*) *attP* sites using the Gateway BP Clonase II Enzyme mix (Thermo Fisher), either directly, or following overlap extension PCR, for the deletion mutants. *P. aeruginosa* deletion mutants (Δ*pqsH*, Δ*pqsR*, Δ*lasA*, and Δ*hcnABC*) were generated as previously described (*117*). Briefly, upstream and downstream fragments were assembled with an FRT-site flanked gentamicin resistance cassette using overlap PCR, and the fragment was cloned into the PCR8/GW/TOPO vector (Invitrogen) by TA cloning and then transferred to the pEX18ApGW (*118*) vector using an LR reaction.

Plasmids were confirmed by sequencing and transformed into the *E. coli* S17-1 λ-pir conjugative strain (*119*). Bi-parental conjugations were performed as described previously between *E. coli* donors and *S. maltophilia* or *P. aeruginosa* recipients (*117*). After conjugation (for four hours for *P. aeruginosa* and overnight for *S. maltophilia*), cells were collected in PBS and plated on LB + irgasan + gentamicin plates. For *S. maltophilia*, the integrated plasmid was excised via selection on LB agar (without NaCl) + 10% sucrose plates. For *S. maltophilia*, gentamicin sensitivity was confirmed by plating. For *P. aeruginosa*, after testing for carbenicillin sensitivity indicating the loss of the plasmid, the gentamicin resistance cassette was removed by a subsequent conjugation with an *E. coli* S17-1 λ-pir conjugative strain carrying the pFLP2 plasmid (*120*). Conjugants were selected on LB + irgasan + carbenicillin plates, streaked on LB agar (without NaCl), and tested for gentamicin sensitivity. Mutations were verified by PCR and sequencing.

For multi-copy complementation of the *smf-1* mutant in *S. maltophilia*, a 5kb PCR fragment encoding the genes for the putative *smf* operon consisting of *smf-1*, *smfB*, *smfC*, and *smfD* (*RS03355*, *RS03360*, *RS03365*, and *RS03370*, respectively) and its putative promoter was amplified from K279a genomic DNA and cloned into pBBR1MCS (*63*) using Gibson assembly. The resulting vector (pSmf) was confirmed through whole plasmid sequencing. The empty vector control or pSmf was electroporated into wild type and mutant backgrounds as described (*121*), and cells selected on LB + chloramphenicol plates, with clones confirmed via PCR.

To facilitate epifluorescence microscopy, GFP and mKO fluorescent tags were introduced in single copy on the chromosome at the *att*:Tn7 in several backgrounds of both *S. maltophilia* and *P. aeruginosa* through double electroporation with a pUC18T-mini-Tn7T-Gm backbone and pTNS3 helper plasmid (*121*, *122*). To facilitate co-culture enumeration, *P. aeruginosa* PA14 mKO was made gentamicin sensitive through Flp-mediated excision of the gentamicin antibiotic-resistance selection marker. Cells were conjugated with *E. coli* S17-1 λ-pir cells carrying the pFLP2 plasmid and selected on LB + carbenicillin plates. The pFLP2 (*120*) plasmid was selected against by plating on LB agar (without NaCl) + 10% sucrose plates, and gentamicin and carbenicillin sensitivity was confirmed by plating.

### Biochemical fractionation

Ten milliliters of *P. aeruginosa* flask supernatant were frozen and lyophilized before resuspension in 2 mL of filtered water. Resuspended supernatant was then separated using a Superdex 30 10/300 GL size exclusion column (GE Healthcare) with filtered water as the mobile phase at a maximal flow rate of 0.7 mL·min^-1^ for 1.5 column volumes at 2.6 MPa for a total of 36.5 mL, collecting 2 mL fractions for a total of 18 fractions. Next, fractions were frozen, lyophilized, and resuspended at a 10x concentration in 1 mL of water. For each fraction, 500 µL was used to test for induction of aggregation, as described above.

### Crystal violet biofilm formation assay

*S. maltophilia* was grown overnight in M63 medium. An overnight culture was diluted to a starting OD600 of 0.05 in fresh medium and added to 8 replicate wells of a 96-well plate. Biofilms were allowed to form in static conditions for 24 h at 37°C and assessed for biofilm formation using crystal violet staining as previously described (*123*).

### Swim motility assay

Swim plates were made with M63 + 0.3% agar. The M63 supplements described above (glucose, ACGU solution, supplement EZ, biotin, nicotinamide, ferric citrate and magnesium sulfate) were added after autoclaving. Plates were allowed to stand at room temperature for 4 h prior to inoculating into the middle of the agar with a dipped pipette tip. Plates were incubated at 37°C for 48 h before imaging on a ChemiDoc (Bio-rad).

### Overlay and invasion assays on semi-solid agar plates

For the overlay assays, overnight cultures of *S. maltophilia* were diluted to an OD_600_ of 0.05 in fresh media and grown to an OD_600_ of 0.15. Then, 30 μl were spotted onto the agar surface of a M63 agar plate, prepared as described for the swim motility assays except with 0.5% agar. Next, the spots were allowed to dry for 10 minutes and incubated at 37°C for 4 h. Overnight cultures of *P. aeruginosa* containing a constitutive mKO fluorescent tag were spun down and washed, diluted to an OD_600_ of 0.015, and 30 μL spots were overlaid on the *S. maltophilia* spots. *S. maltophilia* and *P. aeruginosa* overlay co-cultures were grown for an additional 4 to 16 h at 37°C, after which plates were removed, and an agar cut-out of the co-culture growth was made using the top of a P1000 pipette tip. Agar cut-outs were inverted and gently placed onto the surface of a 96-well glass bottom plate containing 5 μl RedDot™2 Far-Red Nuclear Stain (Biotium) to stain dead cells and imaged with epifluorescence microscopy. Co-culture agar overlay assays were performed in duplicate, with one replicate used for imaging and the other for CFU enumeration.

Agar plates used for bacterial invasion assays were similarly prepared with an identical inoculation protocol for *S. maltophilia* spots. After 4 h, 2 μl of overnight cultures of *P. aeruginosa* were spotted adjacent to *S. maltophilia*, and the plate was incubated for an additional 24 h at 37°C. Agar invasion plates were imaged with a G:Box mini 6/9 Multi fluorescence and chemiluminescence imaging system (Syngene).

Survival of *S. maltophilia* in agar overlay and invasion assays was assessed by removing bacterial biomass from the agar surface with a cotton swab and resuspending it in 1 mL PBS. Bacteria were subsequently mechanically dispersed with a 1 mL syringe and 27G 0.5-inch needle, diluted, and enumerated on LB agar plates supplemented with 10 µg·mL^1^ gentamicin to remove intermixed *P. aeruginosa* cells.

## Data availability

The RNA-seq and whole-genome sequencing (WGS) data have been deposited at the NCBI Short Read Archive (SRA) associated with Bioproject PRJNA1257956.

## ACKNOWLEDGMENTS

We would like to acknowledge the Center for Cancer Research (CCR) Genomics Core for RNA-sequencing and whole-genome sequencing (WGS), Ru-Ching Hsia at the CCR Electron Microscopy Laboratory (EML) for the TEM sample preparation and imaging, the CCR Microscopy Core for assistance with image analysis, the Mayo Clinic Metabolomics Core for the untargeted metabolomics, the Cystic Fibrosis Foundation Isolate Core for providing clinical isolates, George O’Toole at the Geisel School of Medicine in Dartmouth College, Dominique Limoli at Indiana University, and Nicholas Cianciotto at Northwestern University for providing strains, and members of the Ramamurthi lab for conceptual and technical assistance with microscopy. We thank Susan Gottesman, Gisela Storz, and Anthony Martini for comments on the manuscript, and members of the Khare, Gottesman, Storz, and Ramamurthi labs for helpful discussion and feedback. This work utilized the computational resources of the NIH High Performance Computing Biowulf Cluster (http://hpc.nih.gov). This work was supported by the Intramural Research Program of the NIH, National Cancer Insitute, Center for Cancer Research. SKL was supported by a Postdoctoral Research Associate Training (PRAT) Fellowship award 1FI2GM150430-01, and TMZ was supported by a PRAT Fellowship award 1FI2GM137843-01, both from the National Institute of General Medical Sciences.

