## Supplemental Figures, and Supplemental Tables 1,4,5 for "*Stenotrophomonas maltophilia* exhibits defensive multicellularity in response to a *Pseudomonas aeruginosa* quorum sensing molecule"

### SUPPLEMENTARY FIGURES

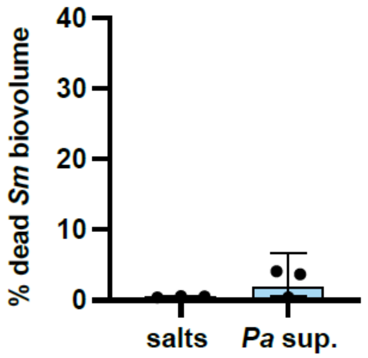

**Supp. Figure 1. *Pa* secreted products do not cause significant *Sm* death.** *Sm* cells were exposed to 50% (v/v) media salts or *Pa* cell-free supernatant for 2 h in static 96-well plates and the biovolume of PI-stained (dead) bacteria was quantified as a percentage of the total biovolume of GFP-tagged bacteria. Data shown are the mean ± SD for three biological replicates. Significance was tested for comparison to the salts control by an unpaired *t*-test (difference was not significant).

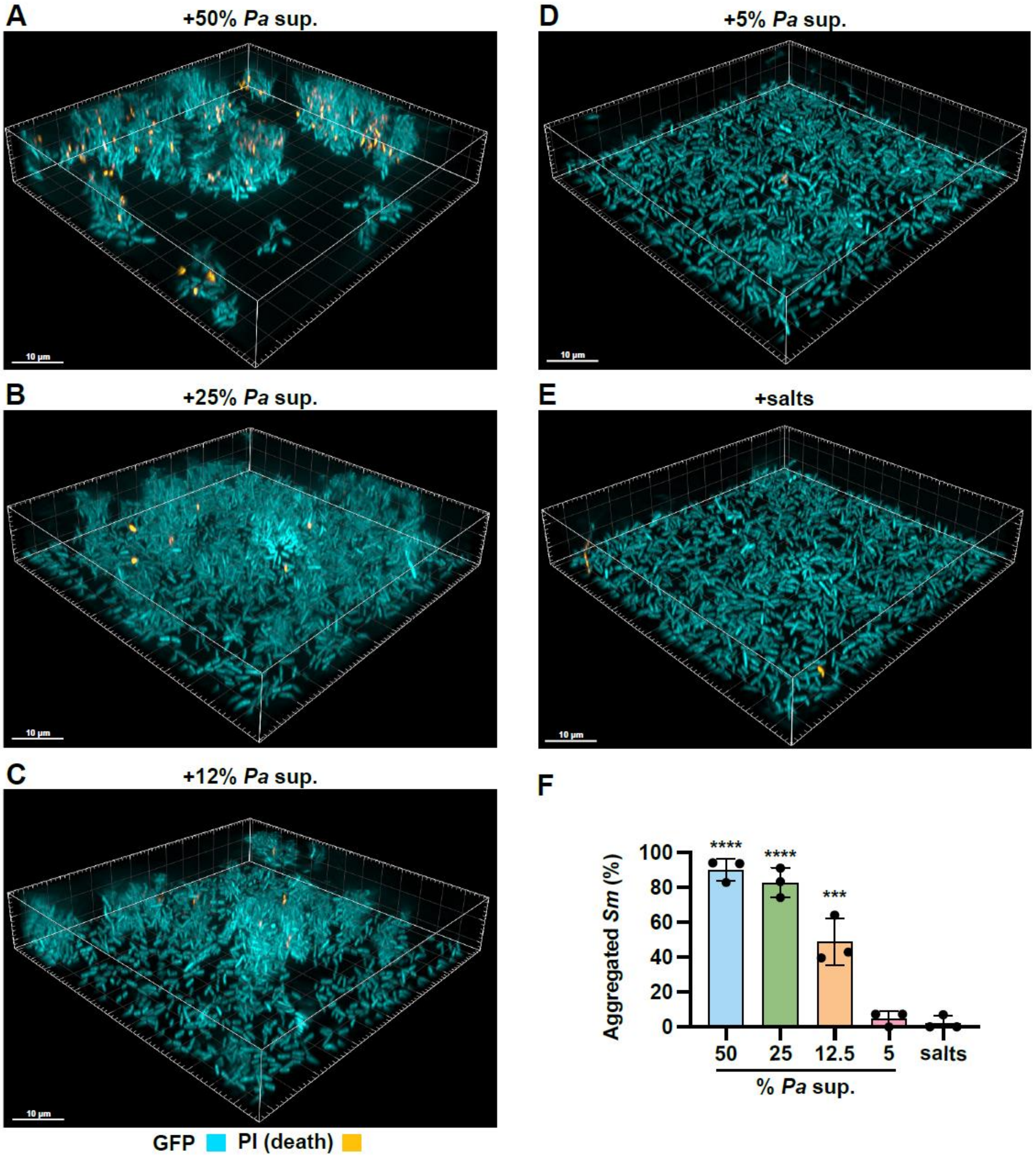

**Supp. Figure 2. *Pa* supernatant-induced *Sm* aggregation is a dose-dependent response. (A-E)** Fluorescent microscopy of GFP tagged and PI-stained *Sm* in static 96-well plates exposed for 2 h to 50, 25, 12.5, or 5% *Pa* cell-free supernatant or salts control, respectively. Representative images of three biological replicates are shown. **(F)** *Sm*

aggregate biovolume derived from representative images of **A-E** and quantified as a percentage of the total *Sm* biovolume. Data shown are the mean ± SD for three biological replicates. Significance is shown for comparison to the salts control, as tested by a one-way ANOVA, followed by the Tukey's test for multiple comparisons (***, $p < 0.001$; ****, $p < 0.0001$).

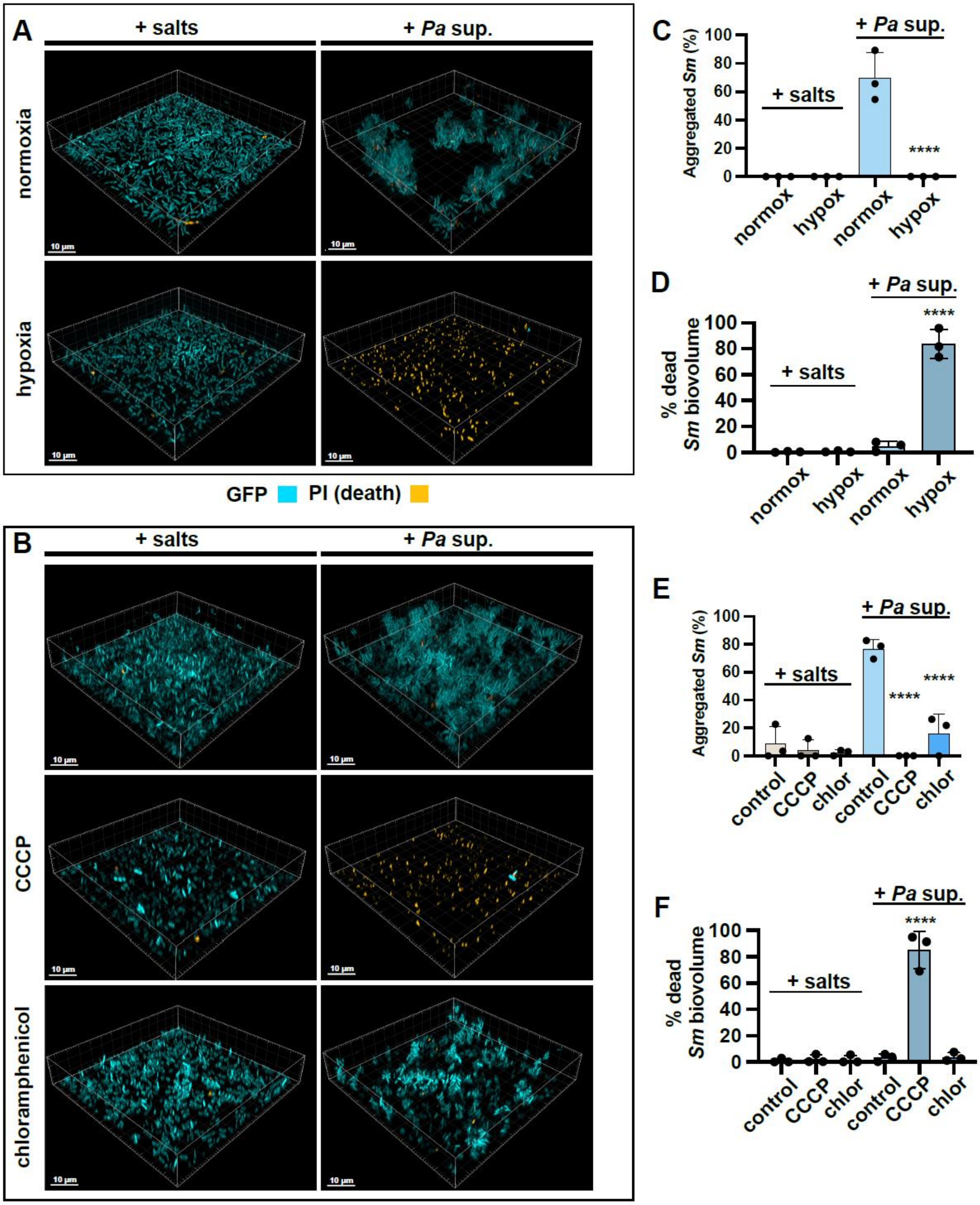

**Supp. Figure 3. *Sm* aggregation in response to *Pa* supernatant is an active behavior**. GFP-tagged *Sm* was stained with PI and pre-treated for 20 minutes with **(A)**

hypoxia and **(B)** CCCP or chloramphenicol and subsequently exposed to *Pa* cell-free supernatant for 2 h and imaged in static 96-well plates. Representative images of three biological replicates are shown. **(C, E)** Biovolume of the resulting *Sm* aggregates was measured as a percentage of the total *Sm* biovolume. **(D, F)** Biovolume of dead *Sm* cells was measured as a percentage of the total *Sm* biovolume. **(C-F)** Data shown are the mean ± SD for three biological replicates. Significance is shown for comparison to **(C, D)** the respective normoxia condition, or **(E, F)** the untreated *Pa* supernatant control, as tested by a one-way ANOVA with the Tukey's test for multiple comparisons (****, $p < 0.0001$).

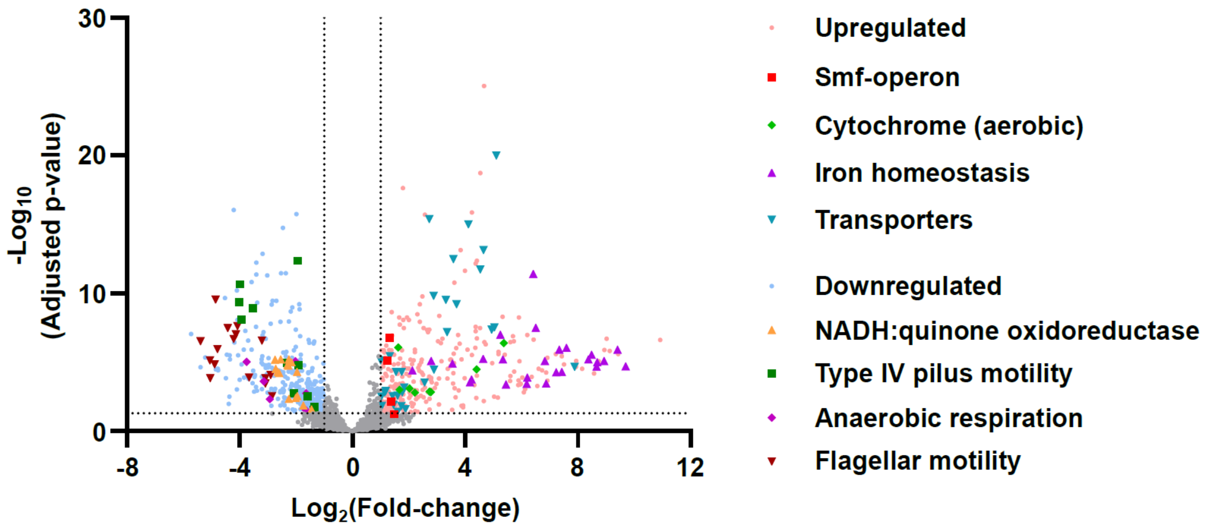

**Supp. Figure 4. *Sm* upregulates the *smf-1* operon and differentially regulates respiration and motility pathways upon exposure to *Pa* supernatant.** WT *Sm* cells were exposed for 30 minutes to either the salts control or *Pa* cell-free supernatant, and transcript levels were measured. Shown are the $\log_{10}$(adjusted *p*-values) and $\log_2$(fold-change).

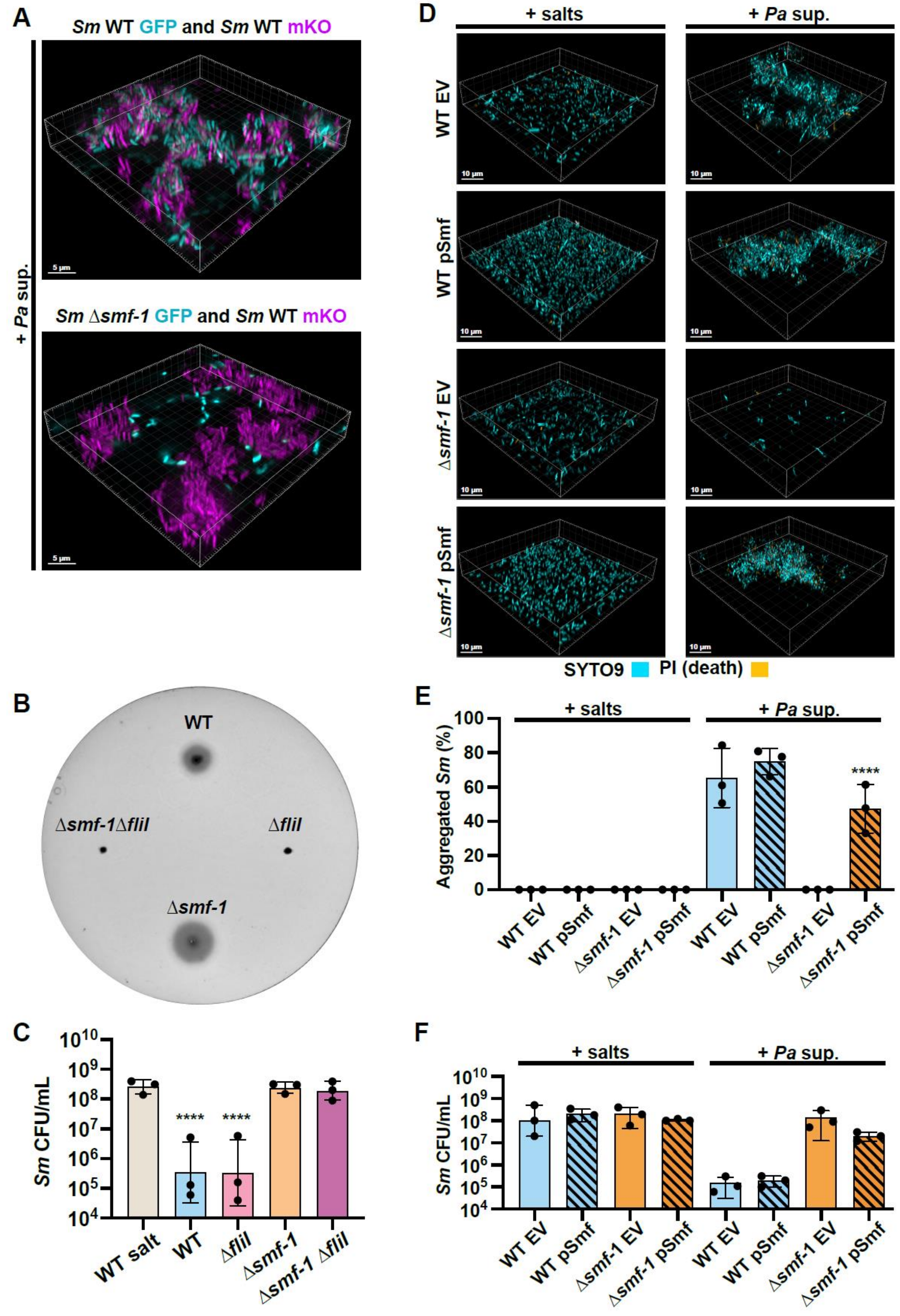
A
Sm WT GFP and Sm WT mKO
Sm Δsmf-1 GFP and Sm WT mKO
+ Pa sup.
5 µm
B
WT
Δsmf-1Δflil
Δflil
Δsmf-1
C
Sm CFU/mL
WT salt
WT
Δflil
Δsmf-1
Δsmf-1 Δflil
****
D
+ salts
+ Pa sup.
WT EV
WT pSmf
Δsmf-1 EV
Δsmf-1 pSmf
10 µm
SYTO9
PI (death)
E
Aggregated Sm (%)
+ salts
+ Pa sup.
WT EV
WT pSmf
Δsmf-1 EV
Δsmf-1 pSmf
****
F
Sm CFU/mL
+ salts
+ Pa sup.
WT EV
WT pSmf
Δsmf-1 EV
Δsmf-1 pSmf

**Supp. Figure 5. *Sm* aggregation requires the Smf-1 fimbriae and is motility-independent. (A)** Microscopy of 1:1 mixtures of WT mKO with either WT GFP or Δ*smf-1* GFP in 96-well plates after 2 h exposure to *Pa* cell-free supernatant. **(B)** Swim motility of WT, Δ*fliI*, Δ*smf-1* and Δ*smf-1* Δ*fliI* on soft agar plates imaged after 24 h. **(C)** CFU enumeration of WT, Δ*fliI*, Δ*smf-1* and Δ*smf-1* Δ*fliI* after 2 h exposure to salts control and *Pa* supernatant. **(D-F)** WT and Δ*smf-1* containing either the empty vector (EV) or one containing the *smf-1* operon (pSmf) after 2 h of exposure to salts and *Pa* supernatant were stained with SYTO9 and PI and assessed via **(D)** microscopy and **(E)** biovolume quantification, or **(F)** CFU enumeration. **(A, B, D)** Representative images of three biological replicates are shown. **(C, E, F)** Data shown are the mean ± SD for three biological replicates. Significance is shown for comparison to **(C)** the WT salts control, or **(E, F)** the EV control, as tested by a one-way ANOVA with the Tukey's test for multiple comparisons (***, $p < 0.001$; ****, $p < 0.0001$).

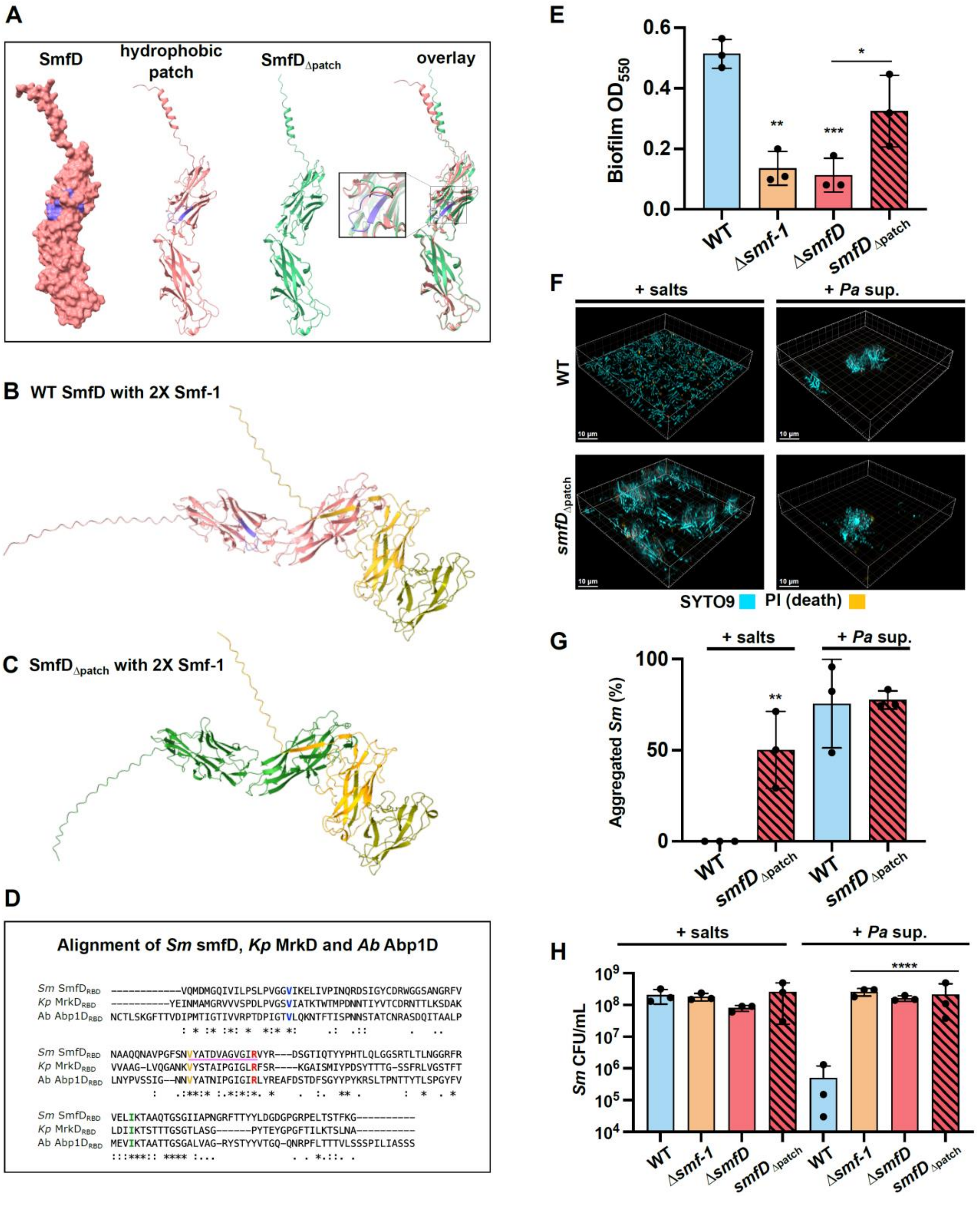

**Supp. Figure 6. SmfD hydrophobic patch alters *Sm* aggregation. (A)** (L to R): SmfD space-filling model, SmfD ribbon drawing with hydrophobic patch depicted in purple,

SmfD hydrophobic patch deletion mutant, and the overlay, with the zoomed-in inset showing the deleted stretch. **(B, C)** Predicted complex model of two Smf-1 subunits, colored in yellow and olive, and **(B)** single WT SmfD or **(C)** single SmfD$_{\Delta patch}$. **(D)** Alignment of receptor binding domains (RBD) of *S. maltophilia* SmfD, *K. pneumoniae* MrkD and *A. baumannii* Abp1D. Residues shown to be required for collagen binding in *K. pneumoniae* MrkD are depicted in blue, green, yellow and red. Region deleted in the *smfD*$_{\Delta patch}$ mutant is underlined in purple and contains two conserved residues from the putative hydrophobic patch. **(E)** Quantification of WT, Δ*smf-1*, Δ*smfD,* and *smfD*$_{\Delta patch}$ biofilm production. **(F, G)** WT, and *smfD*$_{\Delta patch}$ cells were exposed to the salts control and *Pa* supernatant for 2 h in static 96-well plates, stained with SYTO9 and PI, and analyzed via **(F)** microscopy and **(G)** aggregate biovolume. **(H)** CFU enumeration of WT, Δ*smf-1*, Δ*smfD,* and *smfD*$_{\Delta patch}$ cells exposed to the salts control and *Pa* cell-free supernatant for 2 h. **(F)** Representative images of three biological replicates are shown. **(E, G, H)** Data shown are the mean ± SD for three biological replicates. Significance is shown for comparison to the respective WT condition, or the indicated comparison, as tested by a one-way ANOVA with the Tukey's test for multiple comparisons (*, $p < 0.05$; **, $p < 0.01$; ***, $p < 0.001$).

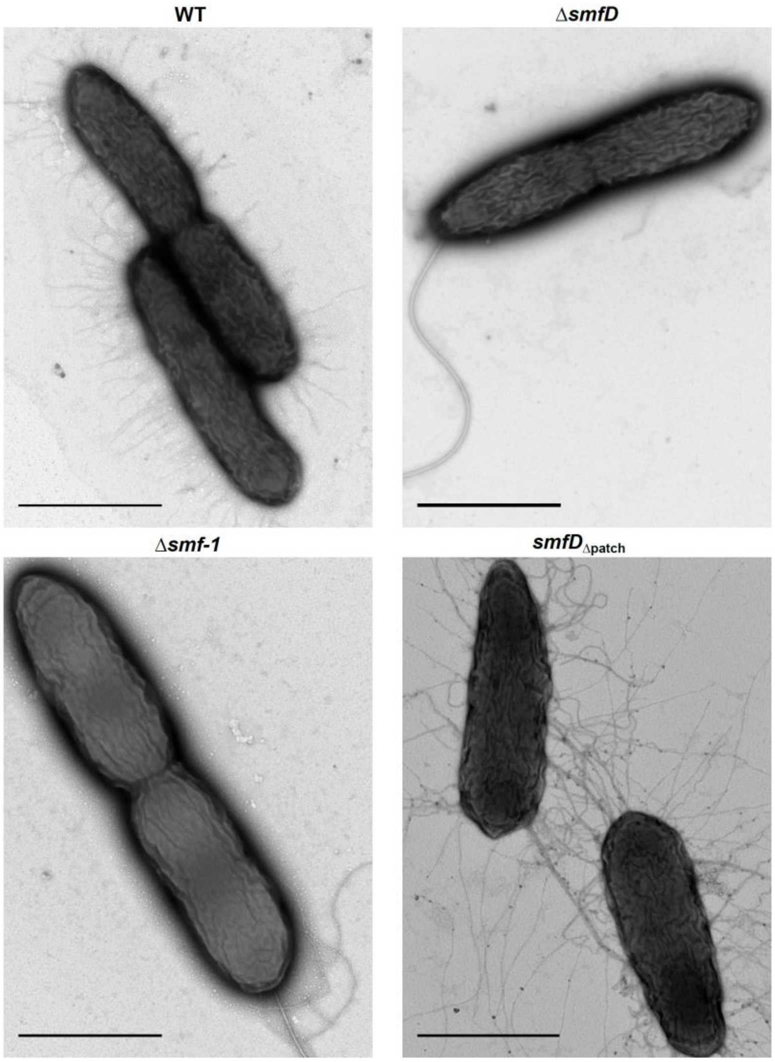

**Supp. Figure 7. The Δ*smf-1* and Δ*smfD* mutants lack fimbriae, while the *smfD*$_{\Delta\text{patch}}$ mutant shows longer, tangled fimbrial structures.** Negative-stained transmission electron micrographs of early log WT, Δ*smf-1*, Δ*smfD,* and *smfD*$_{\Delta\text{patch}}$ mutant cells. Scale bars; 1 μm.

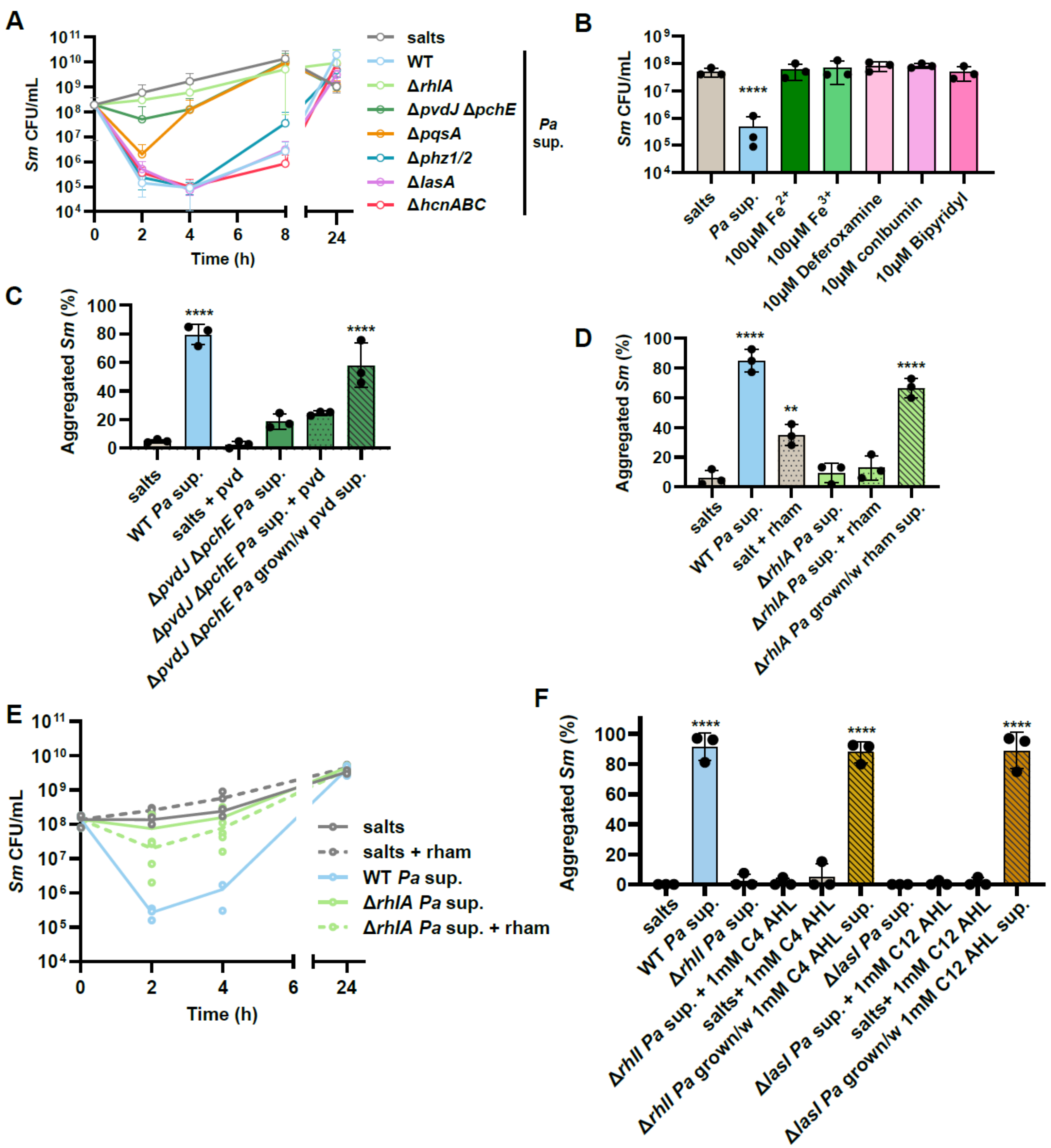

**Supp. Figure 8. *Pa* siderophores, rhamnolipids and N-acyl-homoserine lactone quorum sensing molecules indirectly mediate *Sm* aggregation.** **(A)** CFU enumeration of *Sm* aggregation in response to cell-free supernatant derived from *Pa* WT and mutants deficient for the production of rhamnolipids (Δ*rhlA*), siderophores (Δ*pvdJ*

Δ*pchE*), alkylquinolones (Δ*pqsA*), phenazines (Δ*phz1/2*), the LasA protease (Δ*lasA*) and hydrogen cyanide (Δ*hcnABC*)**. (B)** *Sm* aggregation as measured by CFU enumeration after 2 h exposure to salts, WT *Pa* supernatant, exogenous supplementation of ferrous or ferric iron, or the chelators ferrozine, conalbumin and 2,2 -Bipyridyl**. (C)** Biovolume quantification of *Sm* aggregation in static 96-well plates in response to salts with or without exogenous pyoverdine (pvd), *Pa* WT supernatant, Δ*pvdJ* Δ*pchE* supernatant with or without supplemental pyoverdine, and supernatant from the Δ*pvdJ* Δp*chE* mutant grown with pyoverdine supplementation. **(D)** Biovolume quantification of *Sm* aggregation after 2 h exposure to salts with or without exogenous rhamnolipids (rham), *Pa* WT supernatant, Δ*rhlA* supernatant with or without supplemental rhamnolipids, and supernatant from Δ*rhlA* grown with exogenous rhamnolipids**. (E)** CFU enumeration at the indicated time-points of *Sm* exposed to salts with or without exogenous rhamnolipids, WT *Pa* supernatant, and Δ*rhlA* supernatant with or without exogenous rhamnolipids. **(F)** Biovolume quantification of *Sm* aggregation in response to salts with or without N-butanoyl-L-homoserine lactone (C4 AHL) or N-dodecanoyl-L-homoserine lactone (C12 AHL), *Pa* WT supernatant, Δ*rhlI* supernatant with or without exogenous C4 AHL, supernatant from Δ*rhlI* grown with exogenous C4 AHL, Δ*lasI* supernatant with or without exogenous C12 AHL, and supernatant from Δ*lasI* grown with exogenous C12 AHL. Data shown are the mean ± SD for three biological replicates. Significance is shown for comparison to **(B-D, F)** the salts control, as tested by a one-way ANOVA with the Tukey's test for multiple comparisons (****, $p < 0.0001$). Data for **(B)** was log-transformed prior to statistical testing.

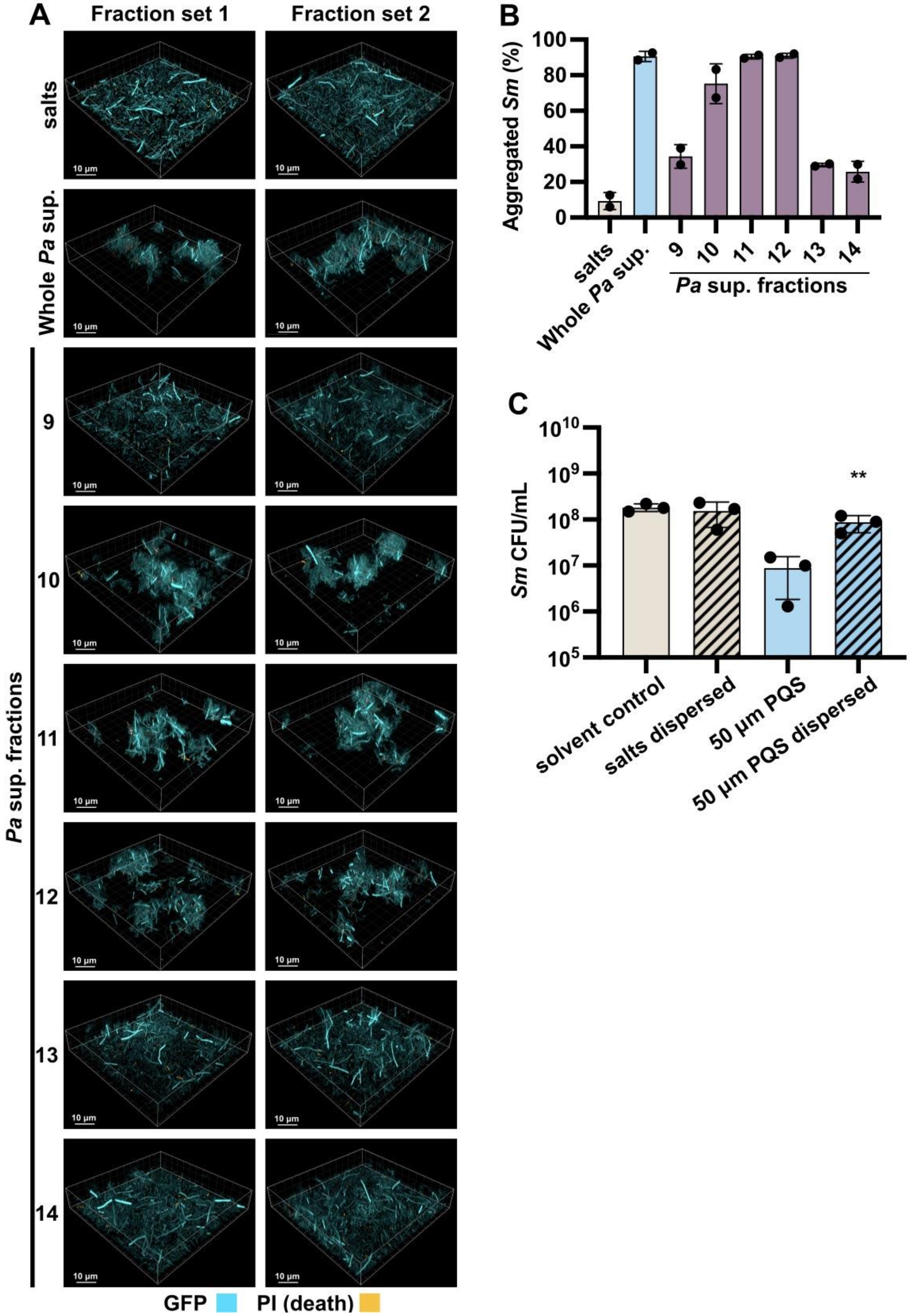
A
Fraction set 1
Fraction set 2
salts
Whole Pa sup.
Pa sup. fractions
9
10
11
12
13
14
10 µm
GFP
PI (death)
B
Aggregated Sm (%)
100
80
60
40
20
0
salts
Whole Pa sup.
9
10
11
12
13
14
Pa sup. fractions
C
Sm CFU/mL
$10^{10}$
$10^{9}$
$10^{8}$
$10^{7}$
$10^{6}$
$10^{5}$
**
solvent control
salts dispersed
50 µm PQS
50 µm PQS dispersed

103

**Supp. Figure 9. Specific *Pa* supernatant fractions and PQS induce *Sm* aggregation.** **(A)** Fluorescent microscopy of GFP tagged and PI-stained *Sm* exposed for 2 h in static 96-well plates to M63 salts medium control, whole *Pa* supernatant, and *Pa* supernatant fractions 9 through 14. Two independent biological replicates (fraction sets) are shown **(B)** Biovolume of the resulting *Sm* aggregates was measured as a percentage of the total *Sm* biovolume. **(C)** CFU enumeration of *Sm* after 2 h of exposure to salts or salts supplemented with 50 µM exogenous PQS followed by mechanical dispersion. Data shown are the mean ± SD for **(B)** two fractions sets and **(C)** three biological replicates. **(C)** Significance is shown for comparison to the respective non-dispersed control, as tested, after log transformation, by a one-way ANOVA with the Tukey's test for multiple comparisons (**, $p < 0.01$).

#### SUPPLEMENTARY TABLES

**Supplementary Table 1. Mutations in evolved *S. maltophilia* isolates.**

| Strains[a] | ORF | Description | Annotation | Position | Mutation |
|---|---|---|---|---|---|
| **Population 1** | | | | | |
| E1.5.1 | SMLT_RS03255 ← \| → *purL* | Chitinase/ Phosphoribosylformylglycinamidine synthase | Intergenic (-324/-489) | 704,573 | C → T |
| | SMLT_RS03350 ← \| → *smf-1* | Hypothetical protein/ Fimbrial protein | Intergenic (-324 / -136) | 734,943 | T → G |
| E1.5.2 | *smf-1* | Fimbrial protein | coding (49/540 nt) | 735,127 | (CGCTCCGC)$_{1\rightarrow 2}$ |
| E1.5.3 | SMLT_RS03350 ← \| → *smf-1* | Hypothetical protein/ Fimbrial protein | Intergenic (-324 / -136) | 734,943 | T → G |
| E1.10.1 | *smf-1* | Fimbrial protein | Coding (293/540 nt) | 735,371 | Δ1 bp |
| | SMLT_RS10875 | Chemotaxis protein CheW | E119E | 2,305,421 | C → T |
| E1.10.2 | SMLT_RS03350 ← \| → *smf-1* | Hypothetical protein/ Fimbrial protein | Intergenic (-324 / -136) | 734,943 | T → G |
| | SMLT_RS14115 ← \| ← SMLT_14120 | Hypothetical protein/ Hypothetical protein | Intergenic (-343/+377; -345/+375) | 3,013,208; 3,013,210 | C → A; C → T |
| E1.10.3 | SMLT_RS03255 ← \| → *purL* | Chitinase/ Phosphoribosylformylglycinamidine synthase | Intergenic (-315/-498) | 704,564 | A → G |
| | SMLT_RS03350 ← \| → *smf-1* | Hypothetical protein/ Fimbrial protein | Intergenic (-324 / -136) | 734,943 | T → G |
| | *hda* → \| ← SMLT_RS05480 | DNA regulatory inactivator/ Nucleotidyltransferase family protein | Intergenic (+366/+133) | 1,174,552 | G → T |
| | *purB* → \| → SMLT_RS15205 | Adenylosuccinate lyase/ cupin domain-containing protein | Intergenic (+416/-240) | 3,235,912 | C → G |
| **Population 2** | | | | | |
| E2.5.1 | SMLT_RS01300 → \| ← SMLT_RS01305 | Sensor histidine kinase/ Zinc-dependent peptidase | Intergenic (+381/+220; +552/+49) | 295,776; 295,947 | C → A; C → A |
| | *smf-1* | Fimbrial protein | coding (344-349/540 nt) | 735,422 | (AGCTGC)$_{2\rightarrow 1}$ |

| | | | | | |
|---|---|---|---|---|---|
| | SMLT_RS09025 | Phage portal protein | V449A | 1,917,865 | A → G |
| E2.5.2 | SMLT_RS03350 ← \| → *smf-1* | Hypothetical protein/ Fimbrial protein | Intergenic (-324 / -136) | 734,943 | T → G |
| | SMLT_RS09025 | Phage portal protein | V449A | 1,917,865 | A → G |
| E2.5.3 | SMLT_RS03350 ← \| → *smf-1* | Hypothetical protein/ Fimbrial protein | Intergenic (-324 / -136) | 734,943 | T → G |
| | *sufD* | Fe-S cluster assembly protein | G115A | 1,198,365 | C → G |
| | SMLT_RS09025 | Phage portal protein | V449A | 1,917,865 | A → G |

[a]First position numeral indicates population 1 or 2; second numeral indicates isolation on 5$^{th}$ or 10$^{th}$ day of passaging; third numeral indicates isolate designation.

120 **Supp. Table 4. Bacterial strains and plasmids used in this study**

| Strain | Description | Source |
|---|---|---|
| *S. maltophilia* | | |
| SB148 | K279a | (*1*) |
| SB490 | K279a GFP | This study |
| SB629 | K279a mkO | This study |
| SB292 | K279a *smf-1*$\Delta_{1bp}$ | This study |
| SB586 | K279a Δ*smf-1* | This study |
| SB630 | K279a Δ*smf-1* GFP | This study |
| *SB589* | K279a Δ*smfD* | This study |
| SB587 | K279a *smfD*$_{\Delta patch}$ | This study |
| SB500 | K279a Δ*fliI* | This study |
| SB501 | K279a Δ*smf-1* Δ*fliI* | This study |
| SB609 | K279a pBBR1MCS EV | This study |
| SB611 | K279a pBBR1MCS pSmf | This study |
| SB610 | K279a Δ*smf-1* pBBR1MCS EV | This study |
| SB615 | K279a Δ*smf-1* pBBR1MCS pSmf | This study |
| AMT0482-08 | *Sm* CF clinical isolate CF077 | CFF Isolate Core |
| AMT0492-03 | *Sm* CF clinical isolate CF082 | CFF Isolate Core |
| AMT0492-08 | *Sm* CF clinical isolate CF087 | CFF Isolate Core |
| AMT0492-12 | *Sm* CF clinical isolate CF091 | CFF Isolate Core |
| *P. aeruginosa* | | |
| 1A3 | PA14 | (*2*) |
| SB489 | PA14 mKO | (*3*) |
| SB92 | PA14 Δ*rhlA* | (*3*) |
| SB459 | PA14 Δ*rhlI* | (*4*) |
| AK625 | PA14 Δ*pvdJ* Δ*pchE* | (*5*) |
| AK619 | PA14 Δ*pqsA* | (*5*) |
| AK652 | PA14 Δ*pqsH* | This study |
| AK660 | PA14 Δ*pqsR* | This study |
| SB521 | PA14 Δ*pqsR* mKO | This study |
| AK681 | PA14 Δ*phz1/2* | (*5*) |
| SB97 | PA14 Δ*lasA* | This study |
| SB460 | PA14 Δ*lasI* | (*4*) |
| SB96 | PA14 Δ*hcnABC* | This study |
| AMT0150-17 | *Pa* CF clinical isolate CF002 | CFF Isolate Core |
| AMT0457-07 | *Pa* CF clinical isolate CF033 | CFF Isolate Core |
| AMT0458-02 | *Pa* CF clinical isolate CF047 | CFF Isolate Core |
| AMT0492-02 | *Pa* CF clinical isolate CF081 | CFF Isolate Core |
| *E. coli* | | |
| DH5α | *E. coli* strain used for cloning | NEB |
| S17-1 λ-pir | *E. coli* strain used for conjugation | (*6*) |
| AK193 | *E. coli* UTI clinical isolate UTI-H | (*7*) |
| AK194 | *E. coli* UTI clinical isolate UTI-P | (*7*) |
| Other species | | |

| | | |
|---|---|---|
| JE2 | USA300_FPR3757 (CA-MRSA)-JE2 | (*8*) |
| SB81 | *Salmonella enterica* | ATCC 29630 |
| SB480 | *Burkholderia cenocepacia*, K56-2 | J. Goldberg |
| SB575 | *Achromobacter xylosoxidans*, CF clinical isolate | D. Limoli |
| SB576 | *Achromobacter xylosoxidans*, CF clinical isolate | D. Limoli |
| SB577 | *Achromobacter xylosoxidans*, ATCC 27061 | D. Limoli |
| **Plasmids** | | |
| pDONRP EX18Gm | Shuttle vector with attP sites and ccdB; Cm$^{r}$ Gent$^{r}$ | (*9*) |
| pEX18ApGW | Gateway-compatible gene replacement vector; Amp$^{R}$ Cm$^{R}$ | (*10*) |
| PCR8/GW/TOPO | Gateway entry vector; Spec$^{R}$ | Invitrogen |
| pFLP2 | FLP recombinase expressing plasmid; Amp$^{R}$ Carb$^{R}$ | (*11*) |
| pSB321 | pDONRPEX18Gm: Δ*smf-1*; Gent$^{r}$ | This study |
| pSB495 | pDONRPEX18Gm: Δ*smfD*; Gent$^{r}$ | This study |
| pSB630 | pDONRPEX18Gm: *smfD*$_{\Delta \text{patch}}$ ; Gent$^{r}$ | This study |
| pSB499 | pDONRPEX18Gm: Δ*fliI*; Gent$^{r}$ | This study |
| pSB601 | pBBR1MCS: empty vector, Cm$^{r}$ | (*12*) |
| pSB614 | pBBR1MCS: pSmf operon complement, *Cm$^{r}$* | This study |
| pAK646 | pEX18ApGW-Δ*pqsH*; Gent$^{r}$ | This study |
| pAK647 | pEX18ApGW-Δ*pqsR*; Gent$^{r}$ | This study |
| pAK959 | pEX18ApGW-Δ*hcn*; Gent$^{r}$ | This study |
| pAK960 | pEX18ApGW-Δ*lasA*; Gent$^{r}$ | This study |

**Supp. Table 5. Primers used in this study**

| Description | Sequence 5′→3′ |
|---|---|
| *smf-1* Δ1bp F | GGGGACAAGTTTGTACAAAAAAGCAGGCTCAAAACACGTCGGCTTACAGGT |
| *smf-1* Δ1bp R | GGGGACCACTTTGTACAAGAAAGCTGGGTAGTATCGGCGGTCTGGTTG |
| *smf-1* Δ1bp. sequencing F | CTGCGCCAGGTCTTCGAG |
| *smf-1* Δ1bp. sequencing R | CCGGAGAAGATCAGTCGCAG |
| Δ*smf-1*, upstream F | GGGGACAAGTTTGTACAAAAAAGCAGGCTCAAAACACGTCGGCTTACAGGT |
| Δ*smf-1*, upstream R | GGGTACGGCTACGATCAGTTCTTGTGCATTCGCTTTTACC |
| Δ*smf-1*, downstream F | GGTAAAAGCGAATGCACAAGAACTGATCGTAGCCGTACCC |
| Δ*smf-1*, downstream R | GGGGACCACTTTGTACAAGAAAGCTGGGTAGTCGTTGATGGTGATGAAGC |
| Δ*smf-1*, sequencing F | CTGCGCCAGGTCTTCGAG |
| Δ*smf-1*, sequencing R | GAGGCGGATGGTGTGTTC |
| Δ*smfD*, upstream F | GGGGACAAGTTTGTACAAAAAAGCAGGCTCACAATACCGTCACCCTCGACC |
| Δ*smfD*, upstream R | CTGGGGCCGCCTTACTCGATCATCGGCATGCTGCCT |
| Δ*smfD*, downstream F | AGGCAGCATGCCGATGATCGAGTAAGGCGGCCCCAG |
| Δ*smfD*, downstream R | GGGGACCACTTTGTACAAGAAAGCTGGGGTTCATTTCCTCCGGCCAC |
| Δ*smfD*, sequencing F | AGCGTGGTCGTGCATCC |
| Δ*smfD*, sequencing R | GACCTGCGTGTAGAGCGC |
| *smfD*$\Delta_{patch}$, upstream F | GGGGACAAGTTTGTACAAAAAAGCAGGCTCACGCTTCGATACCCGCAAG |
| *smfD*$\Delta_{patch}$, upstream R | GGTGCCCGAATCACGATAGACGTTGCTGAAACCCGGGAC |
| *smfD*$\Delta_{patch}$, downstream F | GTCCCGGGTTTCAGCAACGTCTATCGTGATTCGGGCACC |
| *smfD*$\Delta_{patch}$, downstream R | GGGGACAAGTTTGTACAAAAAAGCAGGCTCAGCGGATGCCGATCTTGCT |
| *smfD*$\Delta_{patch}$, sequencing F | GGTGCCCGAGCCCTTGAA |
| *smfD*$\Delta_{patch}$, sequencing R | GGTATTGCCGTGGCCAGC |
| Δ*fliI*, upstream F | GGGGACAAGTTTGTACAAAAAAGCAGGCTCAGCAGATCGAAGGCATCCTCG |
| Δ*fliI*, upstream R | GGCTTAACTCTCTTGTTCCACCAGGTTCATGCATTGGCTCCGGT |
| Δ*fliI*, downstream F | ACCGGAGCCAATGCATGAACCTGGTGGAACAAGAGAGTTAAGCC |
| Δ*fliI*, downstream R | GGGGACCACTTTGTACAAGAAAGCTGGGCGCACCGATATCGTCCAT |
| Δ*fliI*, sequencing F | GACTGCATCCGGATGACATC |
| Δ*fliI*, sequencing R | CAACCGGTCGAGGAAAGC |
| pSmf complementation insert F | GCCCGGGGACACGTTCTCTGGTTCGGTG |
| pSmf complementation insert R | TAGTGGATCCTTACTCGTACTGGATGGTGAAGGTGG |

| | |
|---|---|
| pSmf complementation vector F | GTACGAGTAAGGATCCACTAGTTCTAGAGCGGC |
| pSmf complementation vector R | AACGTGTCCCCGGGCTGCAGGA |
| Δ*pqsH*, A_F | GCTCATCGGTTACCTCTTGCAC |
| Δ*pqsH*, A_R_Gent | TCAGAGCGCTTTTGAAGCTAATTCGAAGAACGGTCATCCGTTGCTC |
| Δ*pqsH*, B_F_Gent | AGGAACTTCAAGATCCCCAATTCGGAGATGGCCGCACAGTAGC |
| Δ*pqsH*, B_R | ACGGTGACTACCACGACCTTG |
| Δ*pqsR*, A_F | CGGATTCTAACCGCATAGGTC |
| Δ*pqsR*, A_R_Gent | TCAGAGCGCTTTTGAAGCTAATTCGAATAGGCATCCCTTATTCCTTTTATTG |
| Δ*pqsR*, B_F_Gent | AGGAACTTCAAGATCCCCAATTCGGCCGCACCAGAGTAGAGC |
| Δ*pqsR*, B_R | CGAGGAAACCCGCAACAAGG |
| Δ*lasA*_A_F | GAGGTGGAAGCCGAGTTTTC |
| Δ*lasA*_A_R_Gm | TCAGAGCGCTTTTGAAGCTAATTCGCATGGGTAGCTCCTGGTC |
| Δ*lasA*_B_F_Gm | AGGAACTTCAAGATCCCCAATTCGTTGTACAACCCCGGCCTG |
| Δ*lasA*_B_R | ACCACCGGCATCATC TTC |
| Δ*hcn*_A_F | CGA GCT TTTCCCCTTCACC |
| Δ*hcn*_A_R_Gm | TCAGAGCGCTTTTGAAGCTAATTCGAAGGTGCATTGCCCTTTC |
| Δ*hcn*_B_F_Gm | AGGAACTTCAAGATCCCCAATTCGTGCTAGGTCCGCGAGGGGTAAATC |
| Δ*hcn*_B_R | CGAGCCACAACTGGTACAGC |
| GentR_F | CGAATTAGCTTCAAAAGCGCTCTGA |
| GentR_R | CGAATTGGGGATCTTGAAGTTCCT |
